# Evidence for reduced immune gene diversity and activity during the evolution of termites

**DOI:** 10.1101/2020.07.09.192013

**Authors:** Shulin He, Thorben Sieksmeyer, Yanli Che, M. Alejandra Esparza Mora, Petr Stiblik, Ronald Banasiak, Mark C. Harrison, Jan Šobotník, Zongqing Wang, Paul R. Johnston, Dino P. McMahon

## Abstract

The evolution of biological complexity is associated with the emergence of bespoke immune systems that maintain and protect organism integrity. Unlike the well studied immunity at the cell and individual level, little is known about the origins of immunity during the transition to eusociality, a major evolutionary transition comparable to the evolution of multicellular organisms from single-celled ancestors. We tackle this by characterizing the immune gene repertoire of 18 cockroach and termite species, spanning the spectrum of solitary, subsocial and eusocial lifestyles. We identified five significant immune gene family contractions and one immune gene family expansion along the spine of a time-calibrated phylogeny, correlating with key transitions in termite sociality. In cross-species comparisons of immune gene expression, we find that termites appear to have evolved a caste-specific social defense system at the expense of individual immune protection. Our study indicates that a major transition in organismal complexity entailed a fundamental reshaping of the immune system optimized for group over individual defense.

## Introduction

The boundaries of individuality have been extended at different stages during the evolution of biological complexity, such as during the evolution of multicellular organisms from single-celled ancestors (Fisher et al., 2013; Michod and Herron, 2006; Pradeu, 2011; Smith and Szathmary, 1997) and the evolution of eusocial animals from solitary ancestors. The fundamental increase of biological complexity has occurred multiple times in the most advanced animal societies, particularly among social insects including some bees, wasps, ants and termites, where the colony has become a dominant unit of selection (Boomsma and Gawne, 2018; Smith and Szathmary, 1997). Immunity is closely tied with these evolutionary transitions, because it is the immune system that defines the boundaries and threats of biological individuality, and is therefore essential for regulating organism integrity (Pradeu, 2011). The evolution of immunity has been well studied at the cell and individual level and efforts to widen understanding to social organisms have been made, such as in bees, thrips, and wasps (Barribeau et al., 2015; Hoggard et al., 2011a; Otani et al., 2016; Turnbull et al., 2010; Turnbull et al., 2012). But a comprehensive exploration of the evolution of immunity has hitherto been lacking during the transition to termite sociality. The termites, along with their nearest living cockroach relatives, represent an excellent system to explore the evolution of immunity due to the presence of a full spectrum of social organization.

Insect immunity has been studied at multiple levels in a small but growing number of insect models. The insect individual immune system has been extensively studied in flies(Hoffmann and Reichhart, 2002; Rolff and Reynolds, 2009) and comprises three principle immune pathways: immune deficiency (IMD), Toll and Janus kinase (JAK)-signal transducer and activator of transcription (STAT). These pathways are typified by pattern recognizing proteins, signaling molecules and effectors, which are responsive to and active against a wide range of insect pathogens. In addition to providing protection at the individual level, social insects have developed a range of group-level social immune traits to protect colonies against infection (Cotter and Kilner, 2010; Cremer et al., 2007; Cremer et al., 2018). Although social immunity has been the focus of much research in a range of social systems in recent years, the evolutionary origins of collective immune defense in social insects has received comparatively little attention. As a combination of traits, ranging from the secretion of antimicrobial compounds to the orchestration of time- and spatially-sensitive collective defenses (Stroeymeyt et al., 2018), the social immune system can be seen to act as a “distributed organ”, much like the conventional immune system of metazoan animals. As with the evolution of the metazoan immune system which is thought to have emerged via the co-option of pre-existing molecular modules and functions into novel defensive pathways, it has been hypothesized that social immune systems originated via similar processes (Pull and McMahon, 2020), with a potentially crucial role for behavioural (Harpur et al., 2019) as well as immune gene adaptations (He et al., 2018; Kutsukake et al., 2019). In line with this view, many genes, including immune-related genes, have been shown to display caste-specific expression patterns (Husseneder and Simms, 2014; Jones et al., 2017; Mitaka et al., 2017; Mitaka et al., 2016; Scharf et al., 2003). In addition, enhanced antimicrobial defenses have been recorded in some social insects compared with their solitary relatives (Hoggard et al., 2011b; Stow et al., 2007; Turnbull et al., 2011; Turnbull et al., 2012).

Immune gene components, particularly immune effectors, have been implicated in the evolution of social immunity. In termites, termicins, a defensin-like gene family, has been duplicated during the evolution of termites (Bulmer and Crozier, 2004), while gram-negative bacteria-binding proteins (GNBP) have acquired a novel fungicidal function in the common ancestor of eusocial termites and subsocial wood roaches via gene duplication (Bulmer et al., 2012), and may play a role in termite collective defensive behavior (Esparza-Mora et al., 2020). In contrast, it has also been hypothesized that sociality could lead to relaxed selection on the individual immune system, potentially via enhanced behaviourally-mediated protection (e.g. via grooming) and reduced pathogen exposure inside colonies. For example, honey bees are typified by a reduction in immune gene diversity (Evans et al., 2006), but this immune gene depletion seems to have preceded the evolution of eusociality in bees (Barribeau et al., 2015), indicating that changes in underlying immune gene evolution are unrelated to group-level defensive or hygienic adaptations linked to sociality in this group.

Comparative analyses of immune gene evolution across a range of independent social groups are required in order to begin navigating across these hypotheses. Along with the intensively studied hymenopteran bee, wasp and ant societies, termites represent an especially important comparative group in this endeavor due to their ancient and evolutionarily distinct origin of sociality, as well as the presence of extant representatives of the full spectrum of social complexity, including some of the most advanced and ecologically successful societies found on earth (Bignell and Eggleton, 2000). Termites possess a rich array of adaptive social immune traits (Rosengaus et al., 2010), which serve to effectively prevent the spread of infectious diseases within colonies (Chouvenc et al., 2012). Genomic analyses of immune gene diversity in a select number of termites and indicate that they may also possess a full complement of canonical insect immune gene pathways(Terrapon et al., 2014), but a comprehensive analysis of total immune gene family evolution across the full spectrum of termite sociality has hitherto been lacking.

We exploited a transcriptomic approach to compare the immune gene repertoire of 18 cockroach and termite species, spanning the full spectrum of solitary and social lifestyles (Fig. 1, Fig. 2), including two solitary cockroach species, two species of subsocial *Cryptocercus* wood-feeding cockroaches, which are the closest living relatives of the termites(Inward et al., 2007a), and 14 species of termites selected from a diverse range of evolutionary lineages across the clade. *Cryptocercus* roaches represent a key lineage in any comparative analysis of termite evolution because they possess important transitional traits such as subsociality, a wood diet with associated protist gut symbionts, and developmental similarities with termites (Inward et al., 2007a; Lo and Eggleton, 2010; Nalepa, 2015). The termite species selected include 8 lower termites representing a range of social modes and ecologies and 6 higher termites belonging to Termitidae, a group which is thought to have undergone further transitions in symbiotic and social evolution (Bucek et al., 2019). Following an investigation into immune gene evolution across a termite phylogeny, we carried out comparative gene expression analyses on representative species bordering the social transition in order to gain deeper insight into the structure of termite immunity.

**Fig. 1.**
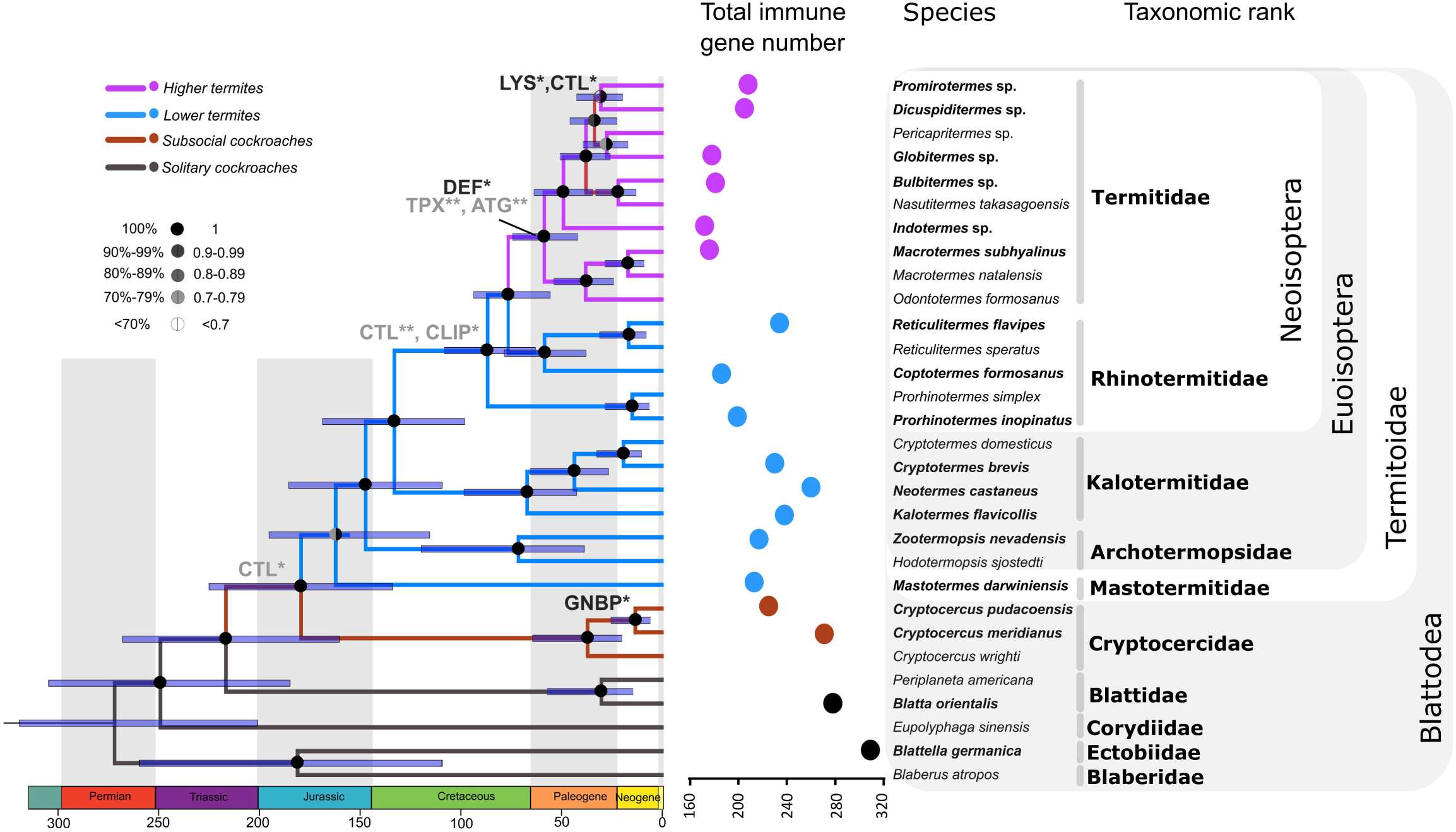
Phylogeny of termites and cockroaches alongside total numbers of identified immune genes. Gene family names in grey and black on the phylogeny indicate significant contractions and expansions of individual gene families, respectively. The gene family evolution analysis was conducted in CAFE. Significance levels of 0.05 (*) and 0.01 (**) are shown.

**Fig. 2.**
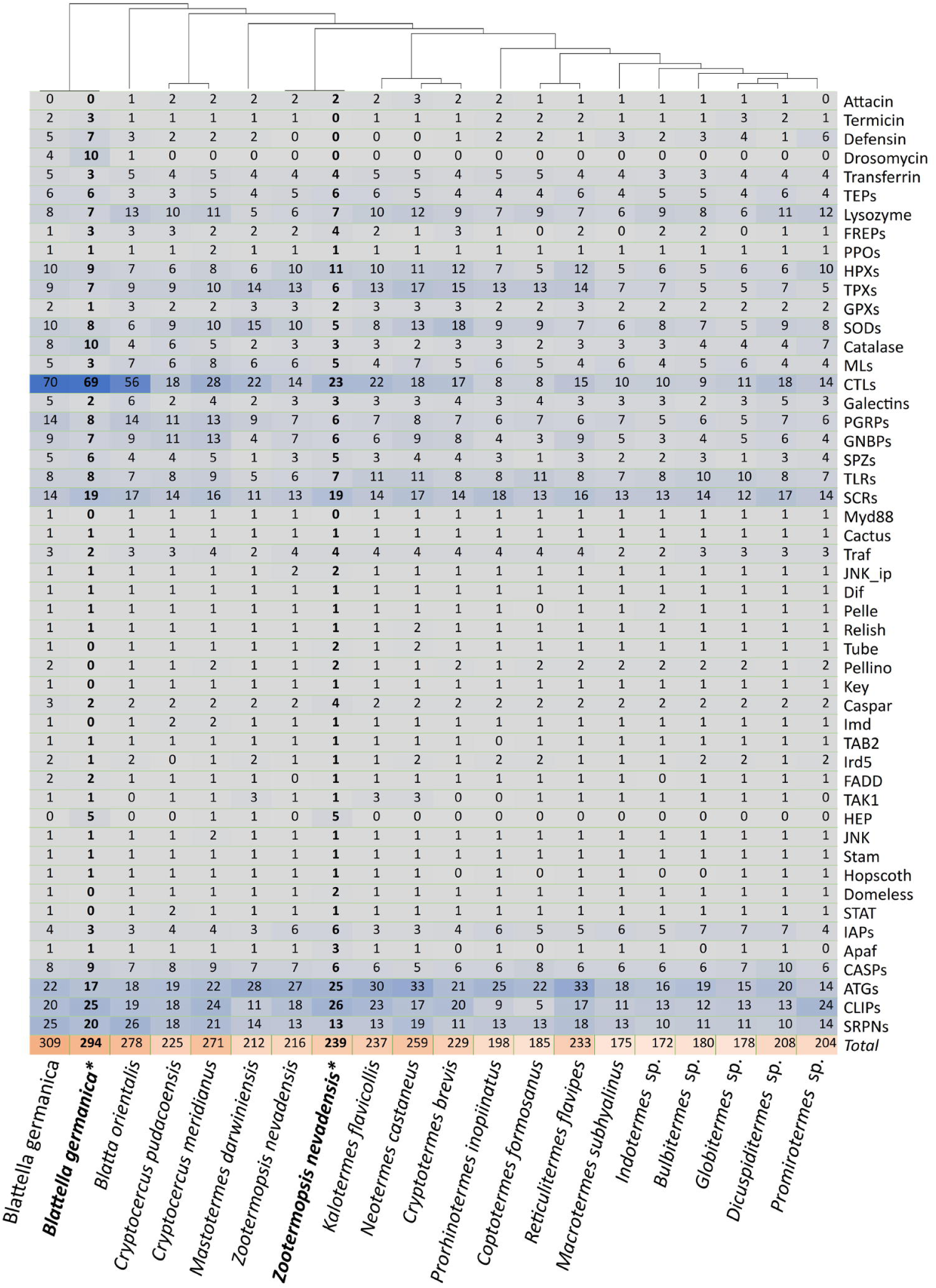
Predicted gene numbers in 50 immune gene families from 18 termite and cockroach species. *: Gene sets of sequenced genomes were used to verify immune gene predictions from our *de novo* transcriptomic data. Columns in bold indicate the number of immune genes estimated from gene sets derived from sequenced genome.

## Results

### Contractions and expansions of immune gene families in termites

We analyzed immune gene evolution over a well-supported termite phylogeny that we reconstructed from 152 single copy orthologs (22898 amino acid positions) of 30 cockroach and termite taxa. Following transcriptome assembly, predicted immune-related genes from 50 gene families were categorized as either receptors, effectors or signaling molecules. Using a combination of identification via hmmsearch and trinotate annotations, we found that each gene family was represented in every cockroach and termite species (Fig. 2), with the noticeable exception of drosomycin, a family of effectors that we find to be lost in termites and wood roaches. An average of 293, 248 and 208 immune-related genes were identified in solitary cockroaches, subsocial wood roaches and social termites, respectively. In a phylogenetic signal analysis, we detected a strong pattern of total immune gene diversity loss during the evolution of termites (Cmean = 0.449, p-value=0.002; Moran’s I=0.055, p-value=0.023; K=1.391, p-value=0.002; K*=0.869, p-value=0.008; λ=0.830, p-value=0.008) with significant positive autocorrelation among species (Fig. S2), particularly among immune gene effector and receptor families (Fig. 2, Fig. S3). For example, C-type Lectins (CTL), peptidoglycan recognition proteins (PGRPs), and attacin genes were notably reduced in number in the majority of termites (Fig. 2). As a control for the potential effect of transcriptome incompleteness, we found no evidence of phylogenetic signal among species for BUSCO scores (Cmean = 0.058, p-value=0.178; Moran’s I= -0.059, p-value=0.467; K=0.371, p-value=0.365; K*=0.489, p-value=0.286; λ<0.0001, p-value=1.0). We then carried out an analysis of gene family evolution by using CAFE to formally test patterns of gene family contraction and expansion over the termite phylogeny. After testing all structures, we found that two λ rates, based on clades with a solitary and sub- or social-system, represented the best fitting model in CAFE (details in Supplementary Text). After applying an error correction, we found the global evolutionary rate of immune gene families in solitary cockroaches (birth/death rate[λ]=0.0037) to be higher than that of subsocial cockroaches and termites (λ=0.0016). Among effector genes, we found that the thioredoxin peroxidase (TPX) gene family had undergone a contraction in the Termitidae crown group (family wide p-value: 0.024, node p-value: 0.0013), while an antimicrobial peptide family, defensin, underwent an expansion in the same group (family wide p-value: 0.011, node p-value: 0.0257) (Fig. 1, Fig. S3). Aside from these immune genes, lysozymes (LYS) also experienced expansions in a node within the higher termites (family wide p-value: 0.02, node p-value: 0.0090). In the receptors, we found that C-type lectins (CTL) underwent two contraction events during the evolution of termite sociality (family wide p-value: 0.011), once in the most recent common ancestor (MRCA) of subsocial wood roaches + social termites (node p-value: 0.0173), and once in the MRCA of Rhinotermitidae + Termitidae (node p-value:0.0021). Interestingly, CTLs appear to have also undergone a re-expansion in higher termites (node p-value: 0.0278), coinciding with the expansion of lysozymes in this node. We also detected evidence of GNBP undergoing an expansion in the common ancestor of subsocial cockroaches (family wide p-value: 0.014, node p-value: 0.0486) and contractions of CLIP (serine protease) in the MRCA of Rhinotermitidae and Termitidae (family wide p-value: 0, node p-value: 0.0403) and autophage related genes (ATG) in the MRCA of Termitidae (family wide p-value: 0, node p-value: 0.0012). Apart from internal nodes shifts, contractions and expansions of gene families were also detected at the tips of termite phylogeny(Fig. S3), including all the gene families mentioned in internal nodes as well as heme-containing peroxidase (HPX) (family wide p-value: 0.005), superoxide dismutase (SOD) (family wide p-value 0.01), and serpin (family wide p-value: 0.005).

### Weak individual immune response in a termite compared with cockroaches

The immune system allows individuals to mount an immune response against microbial infection. To further investigate the evolution of termite immunity, we compared the individual immune responses to infection in a solitary cockroach, *Blatta orientalis*, a subsocial wood-feeding roach, *Cryptocercus meridianus*, and representatives from each caste of the one-piece nesting termite, *Neotermes castaneus*, following direct injection with a cocktail of heat-killed microbes. In the solitary cockroach *B. orientalis*, we found 165 and 263 significantly down- and upregulated genes in immune-challenged individuals respectively (Fig. 3a). Significantly enriched gene ontology (GO) terms of upregulated genes in *B. orientalis* included Toll and PGRP signaling and immune/defense processes (Tab. S1). Among total differentially expressed genes, 25 and 10 represented up- and downregulated immune related genes, respectively (Fig. S4). In the equivalent experiment in the subsocial cockroach *C. meridianus*, we detected a similar pattern to *B. orientalis*, with 248 and 382 genes being significantly downregulated and upregulated, respectively (Fig. 3a). Among the total differentially expressed genes, 24 and 19 represented up- and downregulated immune related genes, respectively (Fig. S4). Overall, solitary and subsocial roaches are characterized by a significant upregulation of immune genes following immune challenge, including members of several gene families ranging from receptor, effector and signaling molecules in both Toll and IMD immune pathways (Fig. 3b, Fig. S4). As in solitary cockroach, PGRP signaling as well as several immune and defense response categories were significantly enriched in upregulated genes in *C. meridianus* (Tab. S2).

**Fig. 3.**
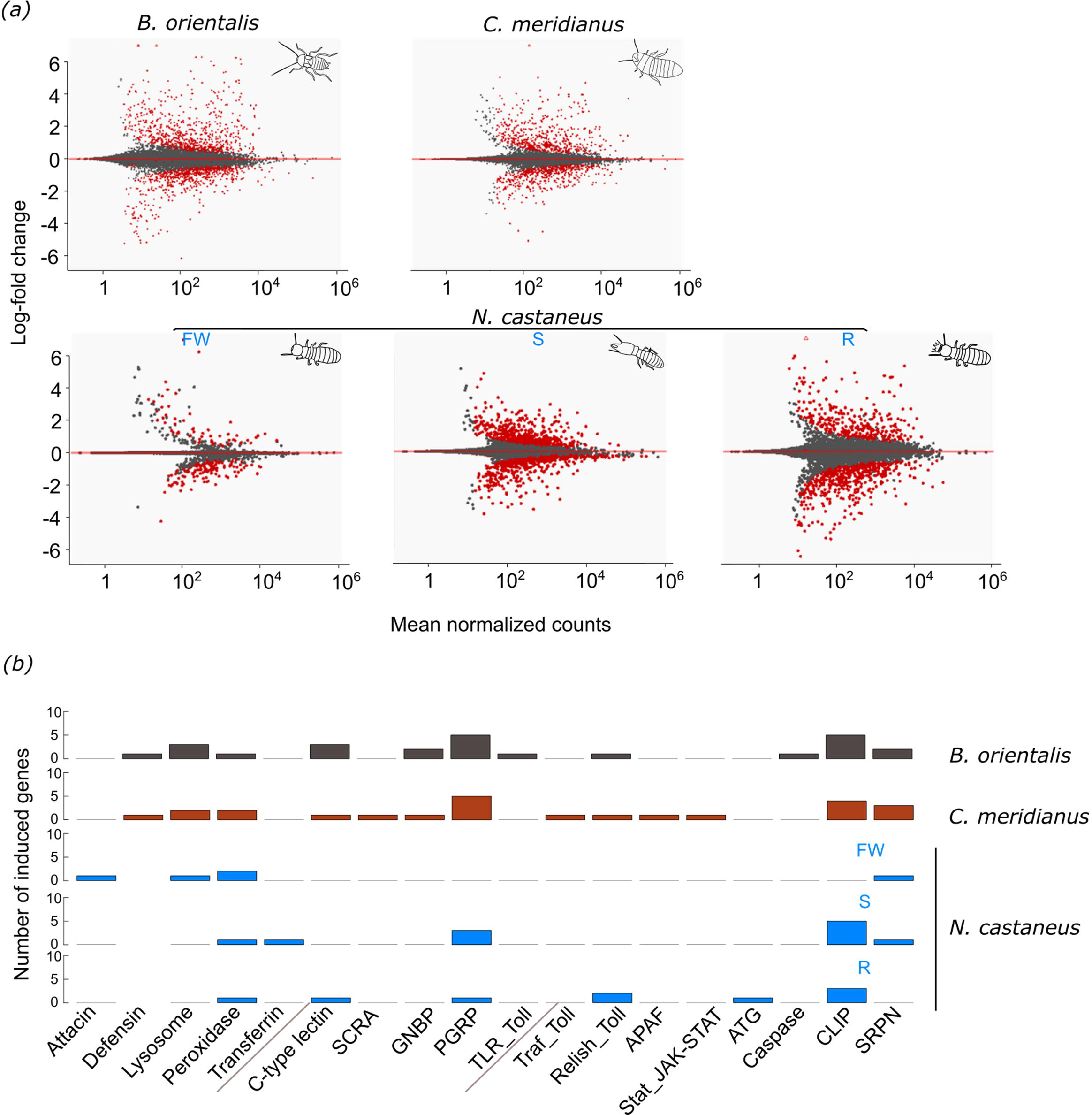
Individual immune response following injection with a cocktail of heat-killed microorganisms versus an equivalent Ringer’s solution. a) Bland-Altman (MA) plots of gene expression in *B. orientalis* and *C.meridianus* (upper panel) or each caste of *N. castaneus*: FW: false workers, S: soldiers, R: reproductives. Red dots in graphs represents differentially expressed genes. b) Cross-species comparison of total number of significantly induced immune genes following experimental injection. Bars in grey, red and blue represent the solitary cockroach, *B. orientalis*, the subsocial cockroach, *C. meridianus*, and the social termite, *N. castaneus*, respectively. FW: false workers, S: soldiers, R: reproductives. Gene families are categorized from left to right into immune effectors, receptors and signaling genes, respectively.

In contrast, a muted response to an equivalent immune challenge was found in termite at the caste-level, with a reduced number of differentially regulated immune genes as well as non-immune genes across all castes, particularly in false workers which upregulated only 30 genes in total in response to treatment (compared with 263 and 382 total upregulated genes in microbe-treated *B. orientalis* and *C. meridianus* individuals versus control, respectively) (log2FoldChange>2, p<0.01) (Fig. 3a). Significantly upregulated genes in false workers (N=30) were not significantly enriched for any GO terms (Tab. S3), while upregulated genes in soldiers (N=161) were significantly enriched in immune-related and transport as well as metabolic process GO terms (Fig. 3a, Tab. S4). Upregulated genes in reproductives (N=220) were significantly enriched in positive regulation of antifungal peptide production and phenol-containing compound biosynthetic processes (Fig. 3a, Tab. S5). Although total upregulated genes in response to immune challenge were higher in soldiers and reproductives compared to false workers, the number of upregulated immune genes was minimal across all castes, with only 9, 11 and 5 immune genes being significantly upregulated in response to immune challenge in *N. castaneus* reproductives, soldiers and false workers, respectively. One immune gene, a HPX was upregulated across all castes, but most upregulated immune genes were caste-specific and functionally non-overlapping, with reproductives and false workers favoring the upregulation of signaling genes and effector molecules (including an Attacin, a Lysozyme and two HPX genes), respectively (Fig. 3b, Fig. S5). Interestingly, however, the number of significantly upregulated unique immune genes in the termite was similar to the number found in solitary and subsocial roach species, when summed across castes (N= 20, 24, and 25 in *N. castaneus, C. meridianus* and *B. orientalis*, respectively) (Fig. S4, S5). Likewise, the number of significantly upregulated unique non-immune genes in the termite was similar to the number found in solitary and subsocial roach species, when summed across castes (N= 313, 358, and 238 in *N. castaneus, C. meridianus* and *B. orientalis*, respectively) (Fig. S6). Of the non-immune genes, only 2 significantly upregulated genes were found to be shared across all three castes. These were a jerky protein homolog-like, and an uncharacterized gene. One gene (poly [ADP-ribose] polymerase 12-like) was significantly upregulated in both false workers and reproductives, while 7 genes were upregulated in both false workers and soldiers and 84 upregulated genes were upregulated in both soldiers and reproductives.

### Caste-specific immunity in the termite *N. castaneus*

We next compared total gene expression differences between castes in the absence of direct immune challenge to understand how caste identity itself shapes constitutive immunity at the individual level. We found that reproductives displayed the highest levels of constitutive immune gene expression, followed by false workers, which can reproduce later in development depending on colony requirements, and then soldiers, which are a permanently sterile terminal caste (Fig. S7). We found that expression of immune related genes could be effectively categorized by caste in a principle component analysis (Fig. 4c). Significantly highly expressed immune genes in reproductives included signaling genes such as Spaetzle, as well as effector molecules Termicin and two Lysozyme genes, while expression of a third Lysozyme, an MD2-like receptor and oxidases were significantly enhanced in false workers. One PGRP gene was significantly highly expressed in soldiers (Fig. 4d). With respect to differentially expressed genes in general, significantly enriched GO terms of highly expressed genes in the reproductive caste included several reproductive and developmental processes as well as pheromone synthesis (Tab. S6), while carboxylic acid biosynthesis was significantly enriched in highly expressed genes of false workers (Tab. S7). No GO terms were significantly enriched in highly expressed genes of soldiers (Tab. S8).

**Fig. 4.**
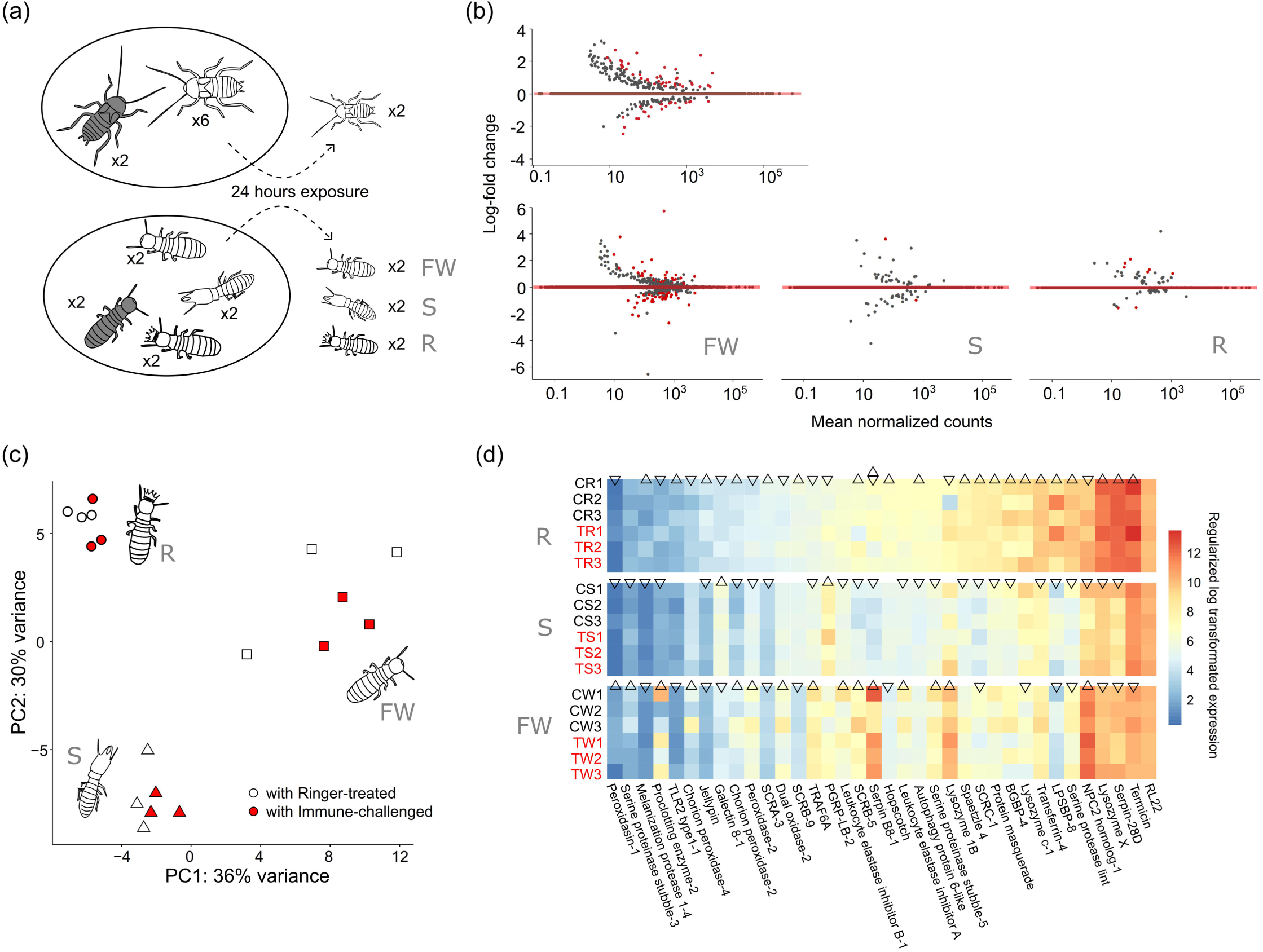
a) A representative diagram of the social group experiment, indicating the design applied to the cockroach *B. orientalis* (upper panel) and the termite *N. castaneus* (lower panel). Individuals marked in grey represent focal individuals challenged by injection with a cocktail of heat-killed microorganisms, or an equivalent Ringer’s control solution. After introduction of injected individuals into social groups, 2 random conspecifics of the injected cockroaches, and all the nestmates of injected termites were sampled for differential gene expression analysis. FW: false workers, S: soldiers, R: reproductives. b) Bland-Altman (MA) plots of gene expression in *B. orientalis* conspecifics (upper panel) or each caste of *N. castaneus* nestmates following exposure to treated focal individuals (lower panel, from left to right: FW: false workers, S: soldiers, R: reproductives). Red dots in graphs represents the differentially expressed genes. c) Principle component analysis (PCA) of total immune gene expression across all three castes of *N. castaneus* from the social experiment, with points in red indicating social groups exposed to immune-challenged focal individuals. d) A heatmap of differentially expressed immune genes following pairwise comparisons among castes. The comparisons were conducted in DESeq2. Expression levels of genes with up-pointing triangles are significantly higher than genes indicated in down-pointing triangles whereas genes with triangles pointing in the same direction pointing indicate non-significance. Gene marked with both an up- and down-pointing triangle are significantly differentially expressed compared with both other castes, whereas genes lacking a triangle are not significantly differentially expressed compared with both other castes.

### Comparison of termite and cockroach gene expression changes in response to a social immune challenge

Recent data indicate that social insect colonies can dynamically adjust interactions in response to infection (Davis et al., 2018; Pull et al., 2018), and segregate between the source of disease and valuable individuals, such as reproductives (Cremer et al., 2018; Naug and Camazine, 2002; Stroeymeyt et al., 2018), in order to keep group fitness. To explore this concept at a molecular level, we quantified gene expression changes in each caste of *N. castaneus* following colony exposure to immune-challenged nestmates (Fig. 4a), and compared these with gene expression changes in the gregarious cockroach, *B. orientalis* following group exposure to immune-challenged conspecifics. The immune challenged individuals of both species was injected with a cocktail of heat-killed microbes, allowing us to exclude the pathogen itself as a cue for social behavior and focus exclusively on the effect of individual health status on social response ^(Hernández López et al., 2017)^. This enabled us to explore how social caste structure influences the response to a social immune threat. In *N. castaneus* we identified a caste-specific response to social immune challenge, with the following number of differentially regulated genes in each caste (upregulated, downregulated): reproductives (1,1), soldiers (1,0), false workers (12,96). Significantly upregulated genes in false workers were related to metabolic functions and chemoreception, including a fatty acid synthase, a trypsin-like protein and a gustatory and odorant receptor. Downregulated genes included transport-related, oxidation-related and protease related genes (Tab. S9). In the equivalent experiment carried out in *B. orientalis*, we found a smaller number of genes to be significantly upregulated (N=9) and downregulated (N=7) following exposure to immune-challenged conspecifics. Upregulated genes in conspecifics included 2 serine proteases, a trypsin-4, an ankyrin repeat and fibronectin type-III domain-containing protein 1 as well as 5 other uncharacterized genes. Downregulated genes contained a hemolymph lipopolysaccharide-binding protein, a troponin T, a protein obstructor-E and 4 other uncharacterized genes. Upregulated genes were enriched for GO terms linked to serine peptidase and hydrolase activity (Fig S8, Tab. S10), although the role of these genes in cockroach immunity remains unclear.

## Discussion

The relationship between the evolution of complexity and immunity is attracting attention as researchers increasing appreciate the interdependency between biological individuality and immunity (Pradeu, 2011, 2019). The evolution of the most advanced forms of eusociality entailed the emergence of a novel form of biological individual – the “superorganism” (Boomsma and Gawne, 2018) – that just as with the evolution of multicellularity, required the obligate loss of independence of previously replicating entities (Fisher et al., 2013; Michod and Herron, 2006). While our understanding of the evolution of individual immunity has increased considerably in recent years, knowledge about the evolution of immunity in social insects has lagged behind (Pull and McMahon, 2020).

We addressed this gap in knowledge in termites by firstly developing a conservative prediction procedure to investigate immune gene family evolution during the transitions through wood roach subsociality to termite sociality. We detected a full repertoire of immune gene families in all *Cryptocercus* and termite lineages except for the antimicrobial peptide drosomycin. Furthermore, we found that early branching termites underwent significant contractions of a few immune gene families followed by minor re-expansions in selected wood roach and termite lineages.

Our reconstruction of immune gene family evolution over a termite phylogeny revealed the loss of drosomycin in the ancestor of *Cryptocercus* wood roaches and termites. Drosomycin was first identified in *Dorsophila* as an antifungal peptide (Zhang and Zhu, 2009). It is unclear whether this loss is caused by ecological shifts or the appearance of social system, or both. But it is possible that the pleiotropic function of newly evolved fungicidal molecules, like GNBP2(Bulmer et al., 2012), which acts synergistically with the AMP termicin(Velenovsky et al., 2016) may have led to functional redundancy and subsequent loss of drosomycin. Alongside evidence of expanded antioxidant genes in cockroaches (Harrison et al., 2018), our observation of contracted TPX (a type of peroxidase known as peroxiredoxins (Radyuk et al., 2001)) in the MCRA of higher termites suggested an important link between antioxidant processing and termite evolution. In addition, we found that CTL, comprising a large proportion of hemolymph lipopolysaccharide-binding proteins (LPSBP) underwent two significant contractions in the MRCA of *Cryptocercus* and termites as well as in the MRCA of Rhinotermitidae and Termitidae. CTLs play an important role in insect innate immunity and can impact infection outcomes for a range of infectious pathogens as well as regulating host microbiota(Zhu et al., 2020). LPSBPs are significantly expanded in cockroaches (Harrison et al., 2018)and are thought to function as opsonins, by binding the surface molecules of invading microorganisms (Jomori et al., 1990; Jomori and Natori, 1991; Jomori and Natori, 1992). A CTL from the hemolymph of *P. americana* has been shown to possess phenoloxidase activity (Arumugam et al., 2017; Chen et al., 1995). Furthermore, LPSBPs may play a possible function in trapping *Blattabacterium* sp. endosymbionts that have leaked from the fat body into the hemolymph, in addition to functioning in the normal cockroach defence mechanism against foreign microbes (Jomori et al., 1990; Kambhampati et al., 1996). The loss of *Blattabacterium* in the ancestor of Euisoptera (all termites excluding Mastotermitidae) may partially explain the pattern of CTL gene depletion, although the significantly reduced diversity of this gene family in both *Cryptocercus* and *M. darwiniensis* indicates that other factors may also be at play. In bees, immune gene depletion seems to have preceded the evolution of eusociality(Barribeau et al., 2015), indicating that immune gene family evolution in Hymenoptera is unrelated to evolutionary transitions in sociality. Although the pattern of immune gene diversity loss in early branching termites appears to contrast with this finding, the significant expansions of genes, including immune genes, in cockroaches compared to other non-social insects(Harrison et al., 2018; Li et al., 2018) could be interpreted as a relative enhancement of immune gene diversity in the ancestral cockroach clade followed by a return to a more representative level of gene diversity in termites.

Aside from a general pattern of immune gene diversity loss in termites, we were able to detect some evidence of gene family re-expansion in some higher termite lineages, potentially resulting from extreme diet diversification and/or shifts in nesting ecology(Donovan et al., 2000). The microbe-enriched lifestyles of which could impose significant selective pressures on immune gene evolution. Equally, modifications to social structure and caste development, or the loss of obligate protist symbionts in the gut could also be major drivers of immune gene evolution in higher termites. However, the extent to each of these large-scale changes in immune family diversity associated with underlying shifts in termite feeding and nesting ecology, microbial symbiosis or sociality requires a more thorough examination in future.

The contractions of immune gene families during termite evolution may reflect a general weakening of individual immunity and/or a specialization of immune responses. We detected similar individual responses to direct immune challenge in the subsocial cockroach *C. meridianus* and the solitary cockroach *B. orientalis*. This suggests that the initial emergence of subsociality was not associated with significant changes to induced immunity. In contrast, a muted individual immune response across all termite castes indicates that the evolution of termite sociality is correlated with a reduced ability to mount a robust immune response. A similar phenomenon has been identified in other social insects including bees and wasps where eusocial insect groups show weaker melanization responses than their close solitary relatives (López-Uribe et al., 2016). This could potentially be the result of trade-offs in selection on individual versus social immunity in more advanced social groups (Cotter and Kilner, 2010).

The social insect colony is a highly organization society with specialized castes. Previous studies in termites have revealed caste specific expression patterns that reflect the specialized functions of castes within colony(Husseneder and Simms, 2014; Jones et al., 2017; Mitaka et al., 2017; Mitaka et al., 2016; Scharf et al., 2003). In this study, we show that constitutive immune gene expression is strongly caste specific in *N. castaneus*, reflecting a division of social roles and indicating a significant degree of caste-specific immune defense. For example, constitutive immune gene expression levels were highest overall in reproductives and lowest in soldiers. A similar finding has been reported in comparisons of workers versus reproductives in bees (Grozinger et al., 2007) and ants ((Graeff et al., 2007), although see (Quque et al., 2019)). Due to the limited number of tested termite species in this study, it is difficult to make generalized statements about common immune gene expression patterns across all termite clades. Nonetheless, our observations clearly reveal a correlation between the evolution of sociality and caste-related immune investment patterns in termites.

Social context plays an important role in coordinating collective behavior in social insects. It has been demonstrated that caste formation can impact immune gene expression in termites (Gao and Thompson, 2015; He et al., 2018). Alongside caste-specific immune gene expression patterns, individuals from different castes may respond to social cues differently, potentially reflecting different levels of investment in individual versus social immunity. Social cues may comprise unique chemical signatures such as cuticle hydrocarbons, which have been shown to be produced by infected worker bees and can evoke an immune response in queens (Hernández López et al., 2017). To investigate this question in termites, we carried out a simplified social challenge experiment whereby the gene expression responses of representatives from each caste of *N. castaneus* were recorded following exposure to immune-challenged false-worker nestmates. For comparison, the equivalent experiment was conducted in a cockroach. Due to the limited material available for the rare subsocial wood roach *Cryptocercus*, we limited our comparisons to *N. canstanus* and the solitary cockroach *B. orientalis*. Interestingly, *B. orientalis* cockroaches exposed to immune-challenged conspecifics had a limited number of differentially regulated genes and no differentially regulated immune- or communication-related genes could be identified. In contrast, termite false workers showed a high number of differentially regulated expressed genes, including upregulated genes in metabolic function- and chemoreception-related activities. Compared with the solitary cockroach, *N. castaneus* appears to be able raise a coordinated caste-specific social immune response, despite it being a single-piece nesting termite species with an intermediate level of social complexity among termites (Inward et al., 2007b). We recorded a negligible impact of social challenge on soldiers and reproductives gene expression indicating that only false workers actively respond to immune-challenged false-worker nestmates, and that they do so by modulating putative sensory and metabolic pathways rather than immune processes. Differentially expressed chemoreception genes in false workers indicate a possible role for chemical communication in coordinating collective social immune responses in *N. castaneus*. However, the importance of behavioural or acoustic cues in termites should be considered as further sources of information in the co-ordination and origins of termite collective defense (Rosengaus et al., 1999). Further work comparing the responses of castes to immune-challenged soldiers and reproductives, in addition to widening study to a greater diversity of termite species, will help to resolve whether the patterns we have observed represent a universal termite response mechanism to social disease challenge.

Using termites as a case study, we have shown that early branching termites underwent significant contractions of major immune gene families followed by minor re-expansions in selected termite lineages. In a cross-species comparison of gene expression, our results reveal a close similarity in induced molecular immunity between solitary and subsocial roach species, despite key ecological, developmental, symbiotic and genomic traits shared by *Cryptocercus* + Termitoidae (Koshikawa et al., 2008; Lo and Eggleton, 2010; Nalepa, 2015; Ohkuma et al., 2008). In comparison with the roach outgroups, we found that termites displayed a dampened response to direct immune challenge at the caste-level. Yet this effect faded when responses were pooled across castes. We find that termites have evolved caste-specific defenses to social as well as individual immune-challenge, reflecting a potential change in focus away from individual defense towards group-level protection and fitness.

Our study shows that the transition to termite eusociality was linked to a significant reconfiguration of termite immune gene diversity and regulation, revealing how a major transition in evolutionary complexity likely entailed fundamental modifications to immune system organization. This study not only provides new insights into the evolution of eusociality in social animals but also facilitate our understanding of the emergence of biological complexity during a major evolutionary transition.

## Methods

### Insects and microorganisms

Larvae and different castes of 9 termite species were extracted from colonies that were kept in the Federal Institute of Materials Research and Testing (BAM), Berlin, Germany. The termite colonies were fed regularly with pre-decayed birch wood or dry grass. An additional 6 species of higher termites were collected from China and Cameroon. Two subsocial wood roaches, *C. meridianus* and *C. pudacuoensis* were collected from Yunnan, China. The solitary cockroaches, *B. orientalis* and *Blattella germanica*, were kept at 26 °C and 75% relative humidity, and were fed with mixed dog food, apples and carrots *ad libitum* until used in experiments. The details of experimental insects are listed in Supplementary Table S11. A Gram-negative bacterium (*Pseudomonas entomophila*, DSM 28517^*T*^), a Gram-positive bacterium (*Bacillus thuringiensis*, DSM 2046^*T*^) and a yeast (*Saccharomyces cerevisiae*, DSM 1333^*T*^) were stored in BAM and cultivated for use in subsequent immune challenge experiments.

### Sample collection and immune challenge experiments

#### Microorganism preparation

*P. entomophila* and *B. thuringiensis* were grown at 28 °C and 30 °C in nutrient broth, respectively. *S. cerevisiae* were grown at 25 °C in universal yeast medium. A growth curve was made for each microorganism. All microorganisms were freshly cultivated, collected during exponential phase, washed twice with Ringer’s solution, and mixed in equal amounts to form a cocktail with a final concentration of 5 x 10^8^ CFU/ml. The suspension of microorganisms was then heat-killed at 95 °C for 10 minutes for subsequent use in experiments.

#### Samples for immune gene characterization

We generated standardized *de novo* transcriptomes for the identification of immune genes from 19 cockroach and termite species, and subsequent comparative analyses of immune gene evolution over the termite phylogeny. Aside from the wood roaches, all insects were prepared for *de novo* sequencing by snap-freezing freshly collected animals in liquid nitrogen. For the wood roaches, colonies collected in the field were preserved in RNAlater following controlled immune challenge. To enrich the expression of immune genes, cockroach adults were challenged by injecting with 5 x 10^6^ cells of heat-killed microorganisms per gram of weight after being swabbed with ethanol. Cockroach larvae and termites were challenged by piercing the cuticle with a sterile needle that had been dipped in a heat-killed microbial suspension, prepared as described above. The wood roaches were frozen in liquid nitrogen or immersed in RNAlater 24 hours after being challenged. All collected samples were preserved at -70 °C until RNA extraction. Termites from the same caste and treatment (non-challenged or challenged) were mixed together for RNA extraction. Cockroaches were extracted individually except larvae (non-challenged or challenged) which were mixed for RNA extraction. Equal amounts of total RNA from extractions were pooled by species for subsequent library construction.

#### Individual immune challenge experiment

In the first gene expression experiment, responses to a direct immune challenge were compared between the solitary oriental cockroach, *B. orientalis*, the subsocial wood roach, *C. meridianus* and three castes of *N. castaneus* (false workers, soldiers, and reproductive). The termite, *N. castaneus*, is a basal “one-piece” termite (in which a piece of wood serves as both food and nest) and has an intermediate form of social complexity. It possesses a sterile soldier caste, a reproductive caste and so-called false workers, which carry out shared tasks such as proctodeal trophallaxis and allogrooming (Davis et al., 2018; De Bie et al., 2006), while retaining the physiological capacity to develop into reproductive individuals under the right colony conditions ^(Korb and Hartfelder, 2008)^. The subsocial wood feeding, *C. meridianus*, which represents the key lineage of *Cryptocercus* that is important in any comparative analysis of termite evolution because of important transitional traits such as subsociality, a wood diet with associated protist gut symbionts, and developmental similarities with termites (Inward et al., 2007a; Lo and Eggleton, 2010; Nalepa, 2015). The cockroach, *B. orientalis*, is reproductively solitary but behaviourally gregarious. As such, *B. orientalis* represented an appropriate comparative control by possessing inherently social behavior while lacking true eusociality. Individuals (N=16 from one cohort of *B. orientalis*, N=16 from 8 colonies of *C. meridianus*, N=32 of each caste from 16 colonies for *N. castaneus*) were weighed and were injected with 5 x 10^6^ cells of heat-killed microbial cocktail per gram of whole body (N=8 cockroaches; N=16 termites of each caste) or an equivalent volume of Ringer’s solution (N=8 cockroaches; N=16 termites of each caste). Following injection, individuals were kept individually with a piece of filter paper. Termites and *B. orientalis* cockroaches were frozen in liquid nitrogen 24 hours following injection, while wood roaches were immersed in RNAlater and stored at -20 °C until transportation. All samples were preserved at -70 °C until RNA extraction.

For sequencing, equal amounts of total RNA from 8 and 4 injected individuals were pooled for termites and cockroaches for each treatment for library preparation, respectively. Each caste, species and treatment were represented by 2 libraries (N= 12, 4 and 4 total libraries for *N. castaneus, C. meridianus* and *B. orientalis*, respectively).

#### Social immune challenge experiment

In the second gene expression experiment, the transcriptional responses of conspecifics of the oriental cockroach *B. orientalis*, and nestmates of the termite, *N. castaneus* were quantified and compared, following social exposure to related individuals challenged with heat-killed microorganisms or an equivalent Ringer’s solution. To enable accurate comparison, we maintained cockroach groups (N=12 groups, each group hatched from a different ootheca) or termite mini-colonies (N=12 mini-colonies, each mini-colony derived from a different mature colony) under equivalent conditions. Groups were comprised of 8 adult cockroaches or 8 termites (4 false workers, 2 soldiers and 2 reproductives derived from the same mature colony) housed inside plastic containers adjusted to maintain a similar volume to insect body surface area ratio across species. *N. castaneus* mini-colonies were housed inside boxes containing a piece of wood which was transferred from the original colony from which the termites were also sourced. An equivalently sized shelter was constructed from egg-box carton and placed inside the containers housing the cockroach mini-groups. All termite mini-colonies were allowed to adjust to their new setting condition over a period of 2 weeks and were inspected regularly. Only those found to contain freshly laid eggs were used in the experiment. For the experiment, two false workers from each of the 12 termite mini-colonies and two cockroaches from each of the 12 mini-groups were randomly selected and weighed for immune challenge, before being swabbed with ethanol. Half of the focal pairs were injected with 5 x 10^6^ heat-killed microbes per gram of whole body (N=6 groups/mini-colonies) while the remaining half were injected with an equivalent Ringer’s solution (N=6 groups/mini-colonies). After injection, focal individuals were individually marked with dark-green dye and returned to the group or colony of origin. Every non-marked individual from the termite mini-colonies, or two randomly selected non-marked cockroaches from each group, were frozen in liquid nitrogen 24 hours following the introduction of injected individuals and stored at - 70 °C until RNA extraction.

For sequencing, equal amounts of total RNA from 4 individuals of each caste, treatment and species from 2 mini-colonies or mini-groups were pooled for library preparation. Each caste, treatment and species had 3 libraries (N=18 and 6 pooled libraries for *N. castaneus* and *B. orientalis*, respectively). For the analysis of gene expression differences between *N. castaneus* castes (Results section: “Caste-specific immunity in the termite *N. castaneus*”), gene expression data from libraries derived from both social immune treatments were combined prior to analysis (N=6 replicates per caste).

#### Total RNA extraction and transcriptome sequencing

For immune gene characterization, total RNA was extracted as described above. For the first and second gene expression experiments, total RNA was isolated from individuals for all species. Due to the large body size, adult cockroaches were cut into 4-6 parts for separate extraction, followed by re-pooling. For the extraction itself, samples were suspended in pre-cooled Trizol (Thermo Fisher Scientific) and homogenized twice at 10 s at 2 M/s with a 5-mm steal bead (Qiagen) using a tissue homogenizer (MP Biomedicals). Total RNA was isolated with a chloroform extraction, followed by isopropanol precipitation, according to instructions from Trizol. Extracted total RNA was dissolved in RNA storage solution (Ambion) and then incubated with 2 units of TurboDNase (Ambion) for 30 min at 37 °C, followed by purification with an RNAeasy Mini kit (Qiagen) according to manufacturer’s instructions. Quantity and quality of RNA were determined by Qubit and Bioanalyzer 2100, respectively. Following pooling described in sample collection part, total RNA was used to construct barcoded cDNA libraries using a NEXTflex™ Rapid Directional mRNA-seq kit (Bioo Scientific). Briefly, mRNA was enriched using poly-A beads from total RNA and subsequently fragmentated. First and second-strand cDNA was synthesized and barcoded with NEXTflex™RNA-seq Barcode Adapters. The libraries were sequenced on an Illumina NextSeq500/550 platform at Berlin Center for Genomics in Biodiversity Research (BeGenDiv).

### Phylogenetic analysis

In addition to sequence datasets from this study, we used another 10 publicly available transcriptomic datasets of cockroaches and termite for phylogenetic inference (Supplementary Table S12). The raw reads were cleaned and filtered before assembled by Trinity (version v2.5.1) (Grabherr et al., 2011)with default parameters (Kmer length: 25). Subsequently, the assemblies were filtered and cleaned before translated into proteins by Transdecoder (version 5.0.1) with a minimum length of 60 amino acids. Raw 454 sequence reads were assembled by using Newbler v2.7 (454 Life Sciences/ Roche). The translated protein sets and an official gene set of *Macrotermes natalensis* (http://gigadb.org/dataset/100057) were used for ortholog analysis by OrthoFinder (version v2.0.0) (Emms and Kelly, 2015). We selected 152 ortholog groups and aligned each group with MAFFT(Katoh and Standley, 2013), masked aligments with trimAI v1.2(Capella-Gutiérrez et al., 2009), and concatenated with Phyutility(Smith and Dunn, 2008) to build a matrix. We employed two different approaches to constructing the phylogeny: maximum likelihood with RAxML (v8.2.12) (Stamatakis, 2014) and Bayesian inference with ExaBayes (v1.4.1) (Aberer et al., 2014). To estimate the divergence times for termites, a molecular clock analysis was performed with PhyloBayes (v4.1) (Lartillot and Philippe, 2004) with following age constraints: all cockroaches and Isoptera: 145.5-315.2 mya (representing the age of the root)(Vršanský, 2002), Cryptocercus and Isoptera: 130-235 mya (Krishna et al., 2013), Kalotermitidae and Rhinotermitidae plus Termitidae: 94.3-235 mya(Krishna and Grimaldi, 2003), Termitidae and Coptotermes plus Reticulitermes: 47.8-94.3 mya (Engel et al., 2011), Reticulitermes and Coptotermes: 33.9-94.3 mya (Engel et al., 2007). Further details in phylogeny reference and molecular dating are available in Supplementary text.

### Immune related protein identification and evolutionary analysis

#### Assembly annotation

For annotation, the full raw reads from 19 species sequenced for immune gene characterization were assembled with Trinity using default parameters. Assembly completeness was assessed by Benchmarking Universal Single-Copy Orthologs (BUSCO v2) with the Arthropod BUSCO set from orthoDB (version 9) (Simão et al., 2015). Each assembly (except *Pericapritermes* sp., due to low completeness, Table S13) was queried against the NCBI nr database by using DIAMOND (Buchfink et al., 2015) and the taxonomic classification of each query assembly was performed using the Lowest Common Ancestor algorithm. The assemblies were annotated by following the guidelines of Trinotate (https://trinotate.github.io/). The proteins of each assembly were predicted by using TransDecoder (v5.2.0) (http://transdecoder.github.io) with a minimum length of 60 amino acids. Homology searches, predictions and domain identifications were performed locally and subsequently integrated into SQLite database at an e-value threshold of 1e-03.

#### Immune gene identification

We adopted a conservative approach to identifying immune gene targets from our transcriptomes. This strategy exploits the cluster information provided by Trinity, which is then used to curate the identification of immune genes. The process we developed first uses HMMER to identify proteins using a domain-based search strategy. Following filtering, HMMER searches are complemented with a blast approach within the trinotate suites, and the application of further quality control steps. The steps are described in detail below.

The first step entailed the modification of a previously published method (Sackton et al., 2017) to quantify the presence of domains containing putative immune functions. Specifically, immune gene families from 31 species in the orthoDB database as well as Termicin and Transferrins from Uniprot (insects) were downloaded and used to construct a set of HMM profile-curated alignments based on all protein families. The complete set of predicted proteins (> 60 amino acids in length) from transcriptomes were searched for matches against predicted immune-related HMMs using HMMER 3.1(http://hmmer.org/). Following domain identification, the HMMER output was subjected to stringent filtering to exclude misidentified transcripts: 1) All targets with E-values > 0.001 for the best domain were excluded. 2) Targets with overall E-value greater than 10^−5^ were disregarded. 3) Targets with multiple HMMs were assigned only to the best e-value HMM. Following these filtering steps, predicted proteins were queried using blastp against the immune gene family database. Proteins were only considered for further analysis when they were assigned to the same immune family as the HMM search. Thirdly, as most genes have multiple immune predicted proteins derived from different isoforms, only one representative isoform (that encoded the protein with the highest overall E-value HMM among all the other proteins from that gene) was chosen for each gene based on the trinity output header. This process excluded multiple isoforms of the same gene and reduced the redundancy of each assembly.

The filtered HMMER outputs were then further selected using annotations from trinotate. Putative gene targets were selected when the output of their predicted proteins from the constructed database matched their annotations of blastp in trinotate. Subsequently, targets were removed when their predicted proteins were shorter than 100 amino acids in immune gene families, except antimicrobial peptides.

We applied an additional layer of filtering to separate isoforms from paralogues and potential gene fragments based on the headers of trinity assembly output. Firstly, because it is theoretically possible that different components from the same subcluster represent spliced isoforms of a single gene, we aligned nucleotide sequences and corresponding predicted proteins from each subcluster against one other using MAFFT and excluded sequences that were variable in length but otherwise identical. Secondly, to account for different fragments of the same gene potentially appearing in different subclusters of a single cluster (and being erroneously described as two separate genes), we ran an additional blastx search on all putative subcluster sequences. If more than one subcluster had an identical target in the top 10 entries of a DIAMOND blastx search (and overlapped by less than 9 amino acids – a value determined by the use of a 25 k-mer parameter during transcriptome assembly), only the longest subcluster was retained (this applied to 13 of 404 putative immune gene sequences). These additional measures enabled us to accurately differentiate between spliced isoforms or fragmented gene sequences and true paralogs.

Before using our immune gene predictions in downstream analyses, we confirmed the reliability of our method by subjecting our pipeline to the completed genomes of *B. germanica* and *Zootermopsis nevadensis*. We applied the procedures described above, aside from the isoform filtering steps, to the official gene sets of *B. germanica* and *Z. nevadensis* to verify that the immune genes identified from our RNAseq data corresponded to data originating from completed genomes. We found that the numbers of estimated immune genes from transcriptome and genome-derived datasets were consistent with each other in both *B. germanica* and *Z. nevadensis*, with minor variations in a limited number of gene families being detected (Fig. 2).

#### Evolutionary analysis of immune gene families

We tested the patterns of immune gene evolution over our termite phylogeny using phylosignal(Keck et al., 2016), which is designed to detect the presence of phylogenetic signal in continuous traits among species. We employed the time-calibrated phylogeny derived from the above phylogenetic analysis and tested the phylogenetic signal of two trait values associated with each tip (species): i) total predicted immune genes derived from each assembly and, ii) associated BUSCO scores as a control for the effect of transcriptome assembly quality.

The expansion and contraction of immune gene families (Fig. S3) was predicted using CAFE 4.0 (-p 0.05)(De Bie et al., 2006), which is based on gene family size and a dated phylogenetic tree. The official gene set of *Z. nevadensis* (Terrapon et al., 2014) and the transcriptome-derived assembly of *Z. nevadensis* were used to estimate the distribution of differences (esterror command in CAFE, -diff 9, determined by the largest difference of gene family sizes between transcriptomes and genomes in *Z. nevadensis* and *B. germanica*) between genome- and transcriptome-derived data, in order to account for any potential bias introduced from the *de novo* transcriptomic approach. Subsequently, the estimated error difference was applied to all species in the dataset. As CAFE allows the application of different death/birth rates at different nodes in the phylogeny (using “lambda -t”), we chose to test different rate structures in order to establish the most suitable model. In addition to the potential structures based on clades with different levels of sociality (solitary, subsocial, social) we tested 16 other structure-based node-clustering methods, as previously described (Kapheim et al., 2015). The node-clustering method firstly calculated independent maximum likelihood lambda values for each node by setting the rate of the focal node differently to the remaining background nodes, and clustered the rates with kmeans clustering. In total 25 structures (M1-M25, Supplementary text) were tested and each structure was repeated 5 times to check for convergence. After testing all structures, we found that the model containing two λ rates, based on clades with a solitary and sub- or social system, was the best fitting model. The significance of the chosen model was then determined by genfamily and lhtest commands in CAFE. The birth and death rate (lambda) of the chosen model was estimated, and the gene families with family-wide p-values < 0.05 were reported.

### Differential gene expression analysis

The raw datasets for the social and individual immune challenge experiments were assembled together and annotated according to the procedures as applied in the phylogenetic analysis section above. Transcript expression following immune challenge in both experiments was quantified by using salmon (Patro et al., 2017). We applied a taxonomy classification with the LCA algorithm in DIAMOND to identify non-target sequences, after which the transcripts with queried targets from Metazoan were considered as host genes and used for further analysis. Differential gene expression was analysed using the R package DESeq2 (Love et al., 2014) using the contrast argument to extract comparisons of interest from DESeq models where all groups from a given experiment are run together (e.g. treatments or castes). In the comparison of gene expression between termite castes as well as in the individual immune challenge experiment we considered genes to be significantly differentially expressed when fold changes > 4 and adjusted p-values < 0.01. Because the responses of nestmates in the social immune challenge experiment were potentially subtle, we considered genes to be significantly differentially expressed when fold changes > 2 and adjusted p-values <0.05. Significantly differentially expressed genes were subject to Gene Ontology (GO) enrichment analysis by the R package goseq with an adjusted p-value cut-off of 0.05. The GOs were extracted from the Trinotate annotation. After GO enrichment analysis, the redundancy of enriched GOs was reduced by using REVIGO (Supek et al., 2011). We calculated the number of differentially expressed genes for each immune protein family to compare immune responses across termite castes (*N. castaneus*) and species (*B. orientalis, C. meridianus, N. castaneus*).

## Acknowledgments

we acknowledge the high-performance computing service of the ZEDAT of the Freie Universität Berlin for providing allocations of computing time and the Berlin Center for Genomics in Biodiversity Research (BeGenDiv) for providing transcriptome sequencing assistance. We thank Jens Rolff, Renate Radek, Sophie Armitage for giving enormous advice and comments on the study. We appreciate Michael T. Monaghan for his advice on phylogenetic analysis.

## Author contributions

D. P. M. and P. R. J. conceived this study. S. H. and T. S. conducted the experiments. S. H., P. R. J. and D. P. M. analyzed the data. D. P. M. and S. H. wrote the manuscript. All authors contributed to the writing of the manuscript.

## Data and materials availability

All raw data associated with the study are available under BioProject PRJNA635910. All codes associated with the study are available at the repository https://github.com/EvoEcoImm/TheEvolutionofTermiteImmunity.

## Additional Information

**Supplementary Information** is available for this paper. Correspondence and requests for materials should be addressed to Dino P. McMahon.

