## Supplementary Text for "Evidence for reduced immune gene diversity and activity during the evolution of termites"

#### Phylogenetic analysis

In order to construct a comprehensive phylogeny, we analyzed 30 transcriptomes and genomes, of which 1 termite genome and 10 available raw data sets were included alongside the 19 assemblies from our study (Tab. S11 and Tab. S12). To facilitate phylogenetic inference, we removed raw reads derived from rRNA and mitochondrial DNA in 19 sequenced species using Botwie2(Langmead and Salzberg 2012) with converted indices built from related sequences of cockroaches, termites and protists from NCBI. Retained reads were assembled by Trinity (version v2.5.1) (Grabherr, et al. 2011)with default parameters (Kmer length: 25) and trimmomatic to clean low-quality reads. After assembling, gene expression was quantified by using Kallisto(Bray, et al. 2016) for each assembly. To reduce redundancy, the highest expressed isoform for each gene was selected with a script in Trinity. Redundancy was further reduced in each assembly by CD-HIT-EST(Fu, et al. 2012) implementing a 95% similarity cut-off. The assemblies were further filtered by Botwie2 to remove rRNA and mitochondrial DNA as we had done previously to the raw reads. Subsequently, the final assemblies were translated into proteins by Transdecoder (version 5.0.1) with a minimum length of 60 amino acids. The raw sequence reads were downloaded from the SRA database in NCBI and the details are listed in Supplementary Table S12. For assembling, we applied the same procedures for raw Illumina sequence reads and assembled the Raw 454 sequence reads using Newbler v2.7 (454 Life Sciences/ Roche).The translated protein sets were used for ortholog analysis by OrthoFinder (version v2.0.0), which is an all-to-all and gene length balanced method to find ortholog groups, suitable for transcriptome data(Emms and Kelly 2015). For the ortholog analysis, we also included an official gene set *Macrotermes natalensis* (http://gigadb.org/dataset/100057).

After ortholog prediction, the single ortholog groups that met the following criteria were selected for matrix building. To mitigate taxon representation bias per orthogroup, we selected orthogroups that included at least one representative of each of the following taxa: 1) *Mastotermes*, 2) *Zootermopsis and Hodotermopsis*, 3) Kalotermitidae (*Kalotermes*, *Neotermes*, *Cryptotermes*), 4) *Coptotermes*, 5) *Reticulitermes*, 6) *Prorhinotermes*. The longest sequence from each selected orthogroup was queried against the ncbi nr database using blast to check for bacterial and protist contamination. Subsequently, these orthogroups were aligned using MAFFT(Katoh and Standley 2013) with the L-INS-i alignment algorithm. To minimize alignment ambiguities, each aligned orthogroup was masked by trimAI v1.2(Capella-Gutiérrez, et al. 2009) with the gappyout function. Orthogroups were then concatenated with Phyutility(Smith and Dunn 2008). An amino acid data matrix with an average of 82.06% gene occupancy per species was assembled from predicted orthogroups. The resulting matrix comprised 152 orthogroups with 22898 amino acid positions and 17.00% missing data.

We employed two different approaches to constructing the phylogeny: maximum likelihood with RAxML (v8.2.12) (Stamatakis 2014)and Bayesian inference with ExaBayes (v1.4.1)(Aberer, et al. 2014). In RAxML, 1000 rapid bootstrap replicates were calculated by employing the PROTGAMMAAUTO model. The parsimony random seed (-p) and bootstrap random seed (-x) were set to 12345. For ExaBayes, two runs were performed and each with four chains. The starting seed (-s) was set to 258. Analyses were run until both runs had average standard deviation of split frequencies (asdsf) below 1% for at least $10^6$ generations. The phylogenetic trees obtained from two different methods, RaxML and ExaBayes, have identical topologies (Fig. 1, Fig. S1). Cryptocercidae and Isoptera are sister groups and form a clade that is closely related to Blattidae. Mastotermitidae is the basal family of the termites and comprises a sister lineage to all other groups. Archotermopsidae is located between Mastotermitidae and Kalotermitidae. Kalotermitidae is a monophyletic grouping in the phylogeny. Rhinotermitidae is a polyphyletic group, comprised of the monophyletic Rhinotermitinae, Heterotermitinae (consisting of *Coptotermes* and *Reticulitermes*), and Psammotermitinae (consisting of *Psammotermes*, *Prorhinotermes*, *Termitogeton*) and Stylotermitinae. Termitidae is monophyletic and a sister group to Rhinotermitinae.

To estimate the divergence times for termites, a molecular clock analysis was performed with PhyloBayes (v4.1) (Lartillot and Philippe 2004). The topology of the phylogenetic tree was constrained to the consensus tree obtained from ExaBayes. An uncorrelated relaxed clock model, using uncorrelated gamma multipliers (-ugam), was applied in our analysis under a birth death prior (-bd) with soft bounds (-sb). Four independent chains were run with 5 fossil calibration points. The following age constraints were employed in this study: all cockroaches and Isoptera: 145.5-315.2 mya (representing the age of the root) (Vršanský 2002), Cryptocercus and Isoptera: 130-235 mya (Krishna, et al. 2013), Kalotermitidae and Rhinotermitidae plus Termitidae: 94.3-235 mya (Krishna and Grimaldi 2003), Termitidae and *Coptotermes* plus *Reticulitermes*: 47.8-94.3 mya (Engel, et al. 2011), *Reticulitermes* and *Coptotermes*: 33.9-94.3 mya (Engel, et al. 2007). We assessed burn-in, convergence among runs, and run performance by examining parameter files with the program TRACER v1.6.0 (Suchard, et al. 2018). Each chain was run for over 10000 cycles, sampling posterior rates and dates with an initial burn in of 20%. Posterior estimation of divergence times was computed from the chain with the highest ESS. As illustrated in the time calibrated phylogenetic tree (Fig. 1), the most recent common ancestor (MRCA) of *Cryptocercus* and termites can be dated to the lower Jurassic, 179.436± 24.1544 (133.939-225.204, 95% confidence interval (CI)) million years ago (mya), which diverged from the Blattidae in the upper Triassic, around 216.657±28.6003 (160.664-267.785, 95% CI) mya. The root of termites is estimated to be 155.341±21.3062 (115.826-195.454,95% CI) million years old from the upper Jurassic. The MRCA of the higher termites, Termitidae, is estimated to be around 58.9309±8.74701 (42.2055-74.7823, 95%CI) million years old from the upper Paleocene and diverged from lower termites around 76.6184±10.5918 (55.8417-93.9591, 95%CI) mya in upper Cretaceous. Although the estimated ages in our study are generally older those derived from mitochondrial or phenotypic data(Engel, et al. 2009; Bourguignon, et al. 2015)and a recent phylogenetic study of cockroach evolution(Evangelista, et al. 2019), our date estimates are in line with a multiple-fossil calibration analysis(Ware, et al. 2010)and a comprehensive recent study of termite evolution(Bucek, et al. 2019).

### Expansion and contraction of immune gene families

We sequenced 15 termite, 2 *Cryptocercus*, and an additional 2 cockroach transcriptomes. After quality control and assembling, each assembly per species contained 120- 210 thousand transcripts with 82.7%-97.7% complete BUSCOs (except *Pericapritermes sp.* with 69.0% BUSCO completeness, which was excluded for further analysis) (Tab. S13).

Immune related genes from 50 families were categorized as either receptor, effector or signaling molecules. Using a combined identification of hmmsearch and trinotate annotation, every gene family was represented by each cockroach and termite species (Fig. 2), except drosomycin, a family of effectors that has been lost in termites and wood roaches.

In the phylosignal analysis, we found no evidence of phylogenetic signal among species for BUSCO scores ($C_{mean}$ = 0.058, p-value=0.178; Moran's I= -0.059, p-value=0.467; K=0.371, p-value=0.365; K*=0.489, p-value=0.286; λ<0.0001, p-value=1.0). Conversely, we detected a strong pattern of total immune gene diversity loss during the evolution of termites ($C_{mean}$ = 0.449, p-value=0.002; Moran's I=0.055, p-value=0.023; K=1.391, p-value=0.002; K*=0.869, p-value=0.008; λ=0.830, p-value=0.008) with significant positive autocorrelation among species (Fig S2).

The following structures (M1-M25) were tested in our CAFE analysis and each structure was repeated 5 times to check for convergence:

M1: 5 λs in solitary cockroaches, subsocial cockroaches, lower termites (except *Rhinotermes*), *Rhinotermes*, and higher termites

M2: 4 λs in solitary cockroaches, subsocial cockroaches, lower termites, and higher termites

M3: 3 λs in solitary cockroaches, subsocial cockroaches, all termites

M4: 2 λs in solitary cockroaches, all subsocial cockroaches and termites

M5: 3 λs in solitary cockroaches, subsocial cockroaches and lower termites (except Rhinotermes), Rhinotermes and higher termites

M6: 2 λs in all cockroaches, and all termites

M7: a common global λ in all species

M8: all nodes have different λ rates

M9 – M25: different λ rates based on kmeans clusters (k from 2 to 17)

After testing all structures, we found that two λ rates, based on clades with a solitary and sub- or social system (structure M4), represented the best fitting model. After applying an error correction, we found the global evolutionary rate of immune gene families in solitary cockroaches (birth/death rate[λ]=0.0037) to be higher than that of subsocial cockroaches and termites (λ=0.0016). Among effector genes, we found that the thioredoxin peroxidase (TPX) gene family had undergone a contraction in the Termitidae crown group, while we find an antimicrobial peptide family, defensin, to have undergone an expansion in the same group. Aside from these immune genes, lysozyme (LYS) also showed expansion in an internal node of higher termites (Fig. S3). In the receptors, we found that C-type lectins (CTL) underwent two contraction events during the evolution of termite sociality (Fig. 1, Fig. S3), once in the MRCA of subsocial wood roaches + social termites, and once in the MRCA of Rhinotermitidae + Termitidae. Interestingly, CTLs appear to have also undergone a re-expansion in higher termites, coinciding with the expansion of lysozymes in this group. We also detected evidence of GNBP undergoing an expansion in the common ancestor of the subsocial cockroaches and contractions of CLIP (serine protease) and autophage related genes (ATG) in the MRCA of Rhinotermitidae and Termitidae.

**Individual experiment: transcriptional responses of immune-challenged individuals**

To explore the relationship between immune response and division of labour in termites, we quantified the number of immune-related genes which were differentially expressed in response to a common immune challenge in three termite castes, a subsocial cockroach and a solitary cockroach. As in the first experiment, we compared individuals that were injected with heat-killed microbes or an equivalent Ringer's control solution, treated insects were kept individually in order to investigate the individual immune response.

In the solitary cockroach *Blatta orientalis*, we detected 263 and 165 significantly down- and upregulated genes in immune-challenged individuals respectively. Upregulated genes were again represented by significantly enriched immune related GO terms (Tab. S1). Among total differentially expressed genes, 25 and 10 represented up- and downregulated immune related genes, respectively (Fig. S4). In an equivalent experiment in the subsocial cockroach *C. meridianus*, we found a similar pattern to *B. orientalis* with 248 and 382 genes to be significantly downregulated and upregulated (log2FoldChange>2, $p<0.01$), respectively. The upregulated genes represented a comprehensive immune response and were significantly enriched in immune related GO terms (Tab. S2). Among total differentially expressed genes, 24 and 19 represented up- and downregulated immune related genes, respectively (Fig. S5).

In *N. castaneus*, only 3 genes were found to be significantly upregulated in immune-challenged individuals of all three castes (log2FoldChange>2, p<0.01). These were a Jerky protein homolog-like, a peroxidase and an uncharacterized gene. One gene (poly [ADP-ribose] polymerase 12-like) was significantly upregulated in both false workers and reproductives, while 7 genes were upregulated in both false workers and soldiers (log2FoldChange>2, p<0.01). Interestingly, the two castes representing terminal moults: soldiers and reproductive, shared 84 upregulated genes in immune-challenged individuals (log2FoldChange>2, p<0.01). Significantly upregulated genes in false workers (N=30) were not significantly enriched for any GO terms (Tab. S3), while upregulated genes in soldiers (N=161) were significantly enriched in immune-related and transport as well as metabolic process GO terms (Supplementary Table S4). Upregulated genes in reproductives (N=220) were significantly enriched in positive regulation of antifungal peptide production (GO:0002804) and phenol-containing compound biosynthetic processes (GO:0046189) (Supplementary Table S5). Five, 11 and 9 immune related genes were significantly upregulated in false workers, soldiers and reproductives, respectively (Fig. S5). Among these gene, a peroxidase was significantly upregulated in all castes.

In a comparison across the 3 species, we found that the immune responses of the 2 cockroach species were similarly comprehensive, with evidence of significant upregulation of receptor genes, signalling components, and effectors. By contrast, termite reproductives and soldiers displayed a similar but relatively reduced pattern of immune gene expression following immune-challenge. The response of false workers was comparatively even weaker, with differential expression limited to a serpin and 3 effector genes: attacin, lysozyme and peroxidase (Fig. 3).

**Caste-specific immunity in the termite *N. castaneus***

We also compared overall caste-specific genes expression patterns by grouping treatments and comparing differentially expressed genes between castes. We found genes to be significantly grouped by caste (Fig. S9, Fig. S10, Fig. S11). Compared with both false workers and soldiers, reproductives harboured highly expressed genes that were significantly enriched for reproductive and developmental processes as well as in pheromone synthesis, whereas lowly expressed genes were significantly enriched for oxidative phosphorylation (GO:0006119) (log2FoldChange>2, p<0.01) (Tab. S6). In false workers, highly expressed genes were significantly enriched in carboxylic acid biosynthetic processes (GO:0016053) whereas lowly expressed genes were enriched for multi-multicellular organism processes (GO:0044706) compared to both soldiers and reproductive (Tab. S7). Compared to both false workers and reproductive, soldiers harboured highly expressed genes that were enriched, but not significantly, for muscle related activity and development related GO terms, whereas lowly expressed gene were significantly enriched for developmental processes (GO:0032502), proteolysis (GO:0006508), defecation rhythm (GO:0035882) and macromolecule catabolic processes (GO:0009057) (Tab. S8).

Interestingly, we found that expression of immune related genes could be effectively categorized by caste in a principle component analysis (Fig. 2). Reproductives displayed the highest levels of immune-related gene expression, when compared with both soldiers and false workers (Fig. S7).

**Social experiment: transcriptional responses of nestmates exposed to immune-challenged individuals**

In this experiment, we studied the social immune responses of different *N. castaneus* castes following exposure to immune-challenged or Ringer's solution-injected individuals. In order to record transcriptional responses, we challenged two false workers and then returned them to a mini-colony composed of 6 members, comprised of 2 (non-challenged) individuals of each caste, i.e. 2 false workers, 2 soldiers, and 2 reproductives (king and queen).

We investigated gene expression responses of different castes between treatments, following 24 hours of exposure to 2 immune-challenged or Ringer-injected false workers (Fig 4). Among reproductives, we detected 1 significantly upregulated (Fatty-acid amide hydrolase 2 like) and one significantly downregulated gene (Glucose dehydrogenase [FAD, quinone]) between treatments (log2FoldChange>1, $p<0.05$). Among soldiers, 1 gene was significantly upregulated (Guanylate cyclase 32E like) while no genes were downregulated (log2FoldChange>1, $p<0.05$). By contrast, among false workers, 12 genes were significantly upregulated while 96 genes were significantly downregulated(log2FoldChange>1, $p<0.05$). Upregulated genes included Fatty acid synthase-1, Trypsin-1-like, Thiamine transporter 2, Zinc finger protein like, probable cytochrome P450, and a gustatory and odorant receptor 24-like isoform X2, mite allergen Der f 3-like, and 5 uncharacterized genes. The downregulated genes (Tab. S9) mainly comprised transport-related, oxidation-related and protease related genes, and included 6 immune-related genes (5 serine proteases and 1 Chorion peroxidase).

In the equivalent experiment using the solitary cockroach *B. orientalis*, conspecific individuals between treatments exposed to immune-challenged or Ringer's solution-injected showed significant upregulation and downregulation of 9 and 7 genes, respectively (log2FoldChange>1, $p<0.05$). Upregulated genes in conspecifics included 2 serine proteases, a trypsin-4, an Ankyrin repeat and fibronectin type-III domain-containing protein 1 as well as 5 other uncharacterized genes. Upregulated genes were enriched in the "serine-type peptidase activity" molecular functional (MF) GO term (Tab. S10). Downregulated genes contained a Hemolymph lipopolysaccharide-binding protein, a troponin T, a Protein obstructor-E and 4 other uncharacterized genes.

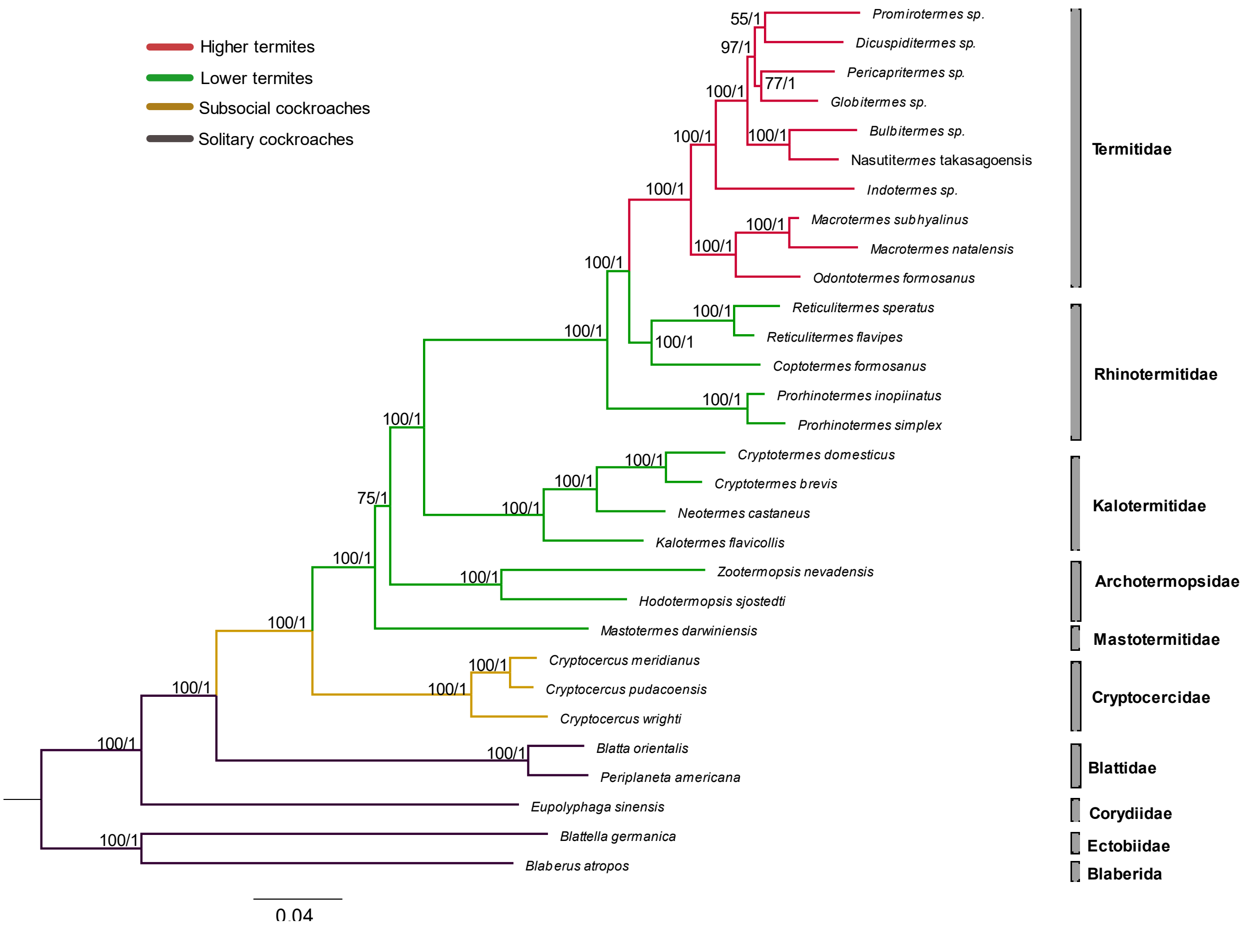


**Figure S1**. Phylogeny of termites based on RAxML and Exabayes. The number on each node represents support of boostrap values from RAxML/likelihood score from Exabayes. Different colors of lines indicate traditional classification of termites and cockroaches.

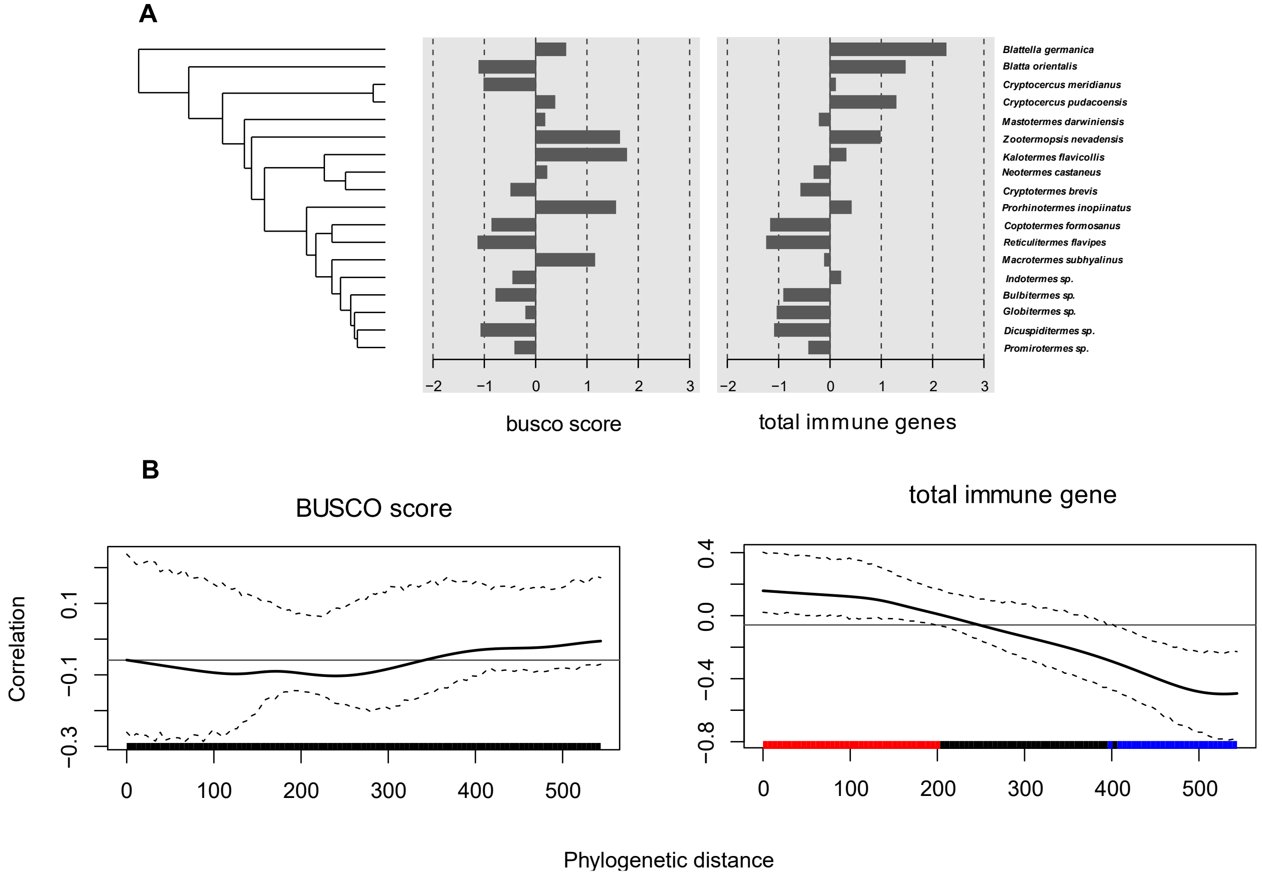


**Figure S2.** Phylosignal analysis. (A) Time-calibrated phylogeny and corresponding phylogenetic signal of two trait values associated with each tip (species). *Left*: associated BUSCO scores as a control for the effect of transcriptome assembly quality. *Right*: total predicted immune genes derived from each assembly. (B) Phylogenetic correlograms displaying the extent of trait autocorrelation among related tips, revealing the pattern and location in the taxonomy of any detected phylogenetic signal. Higher and lower levels of autocorrelation than expected by chance are depicted in red and blue, respectively.

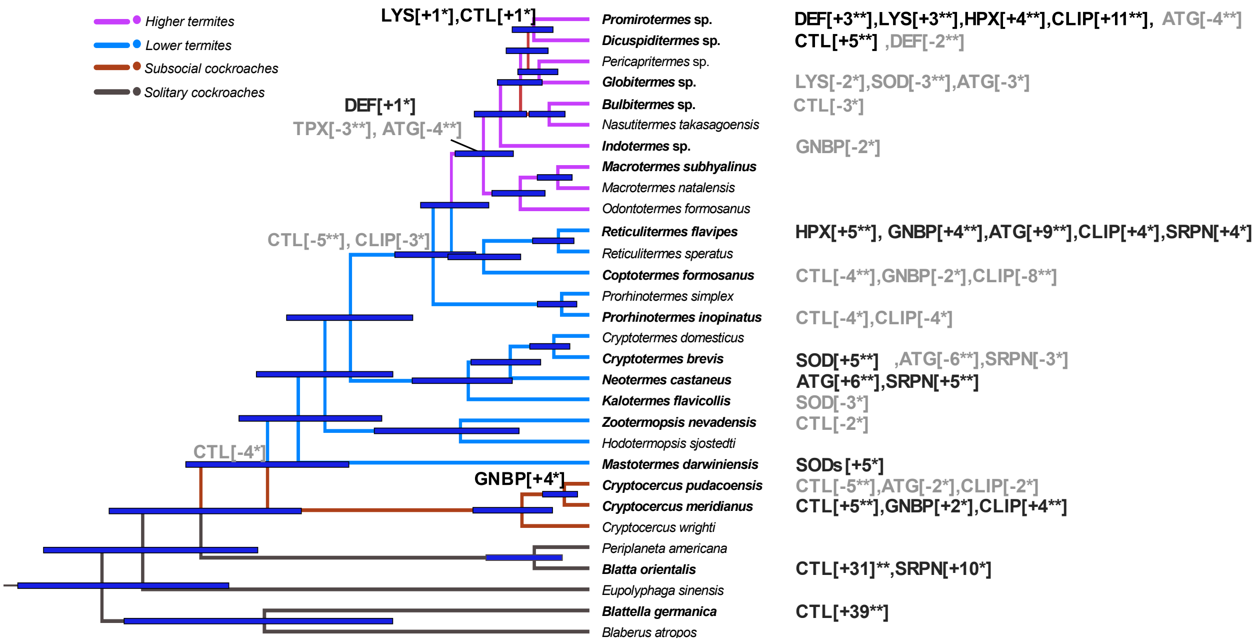


**Figure S3.** Gene family names in grey and black on the phylogeny indicate significant contractions and expansions of individual gene families, respectively. The gene family evolution analysis was conducted in CAFE. Significance levels of 0.05 (*) and 0.01 (**) are shown. The gene number of contraction or expansion in each family were indicated in square brackets. The contractions and expansion in the right panel indicate the significant changes in gene families in individual tested species.

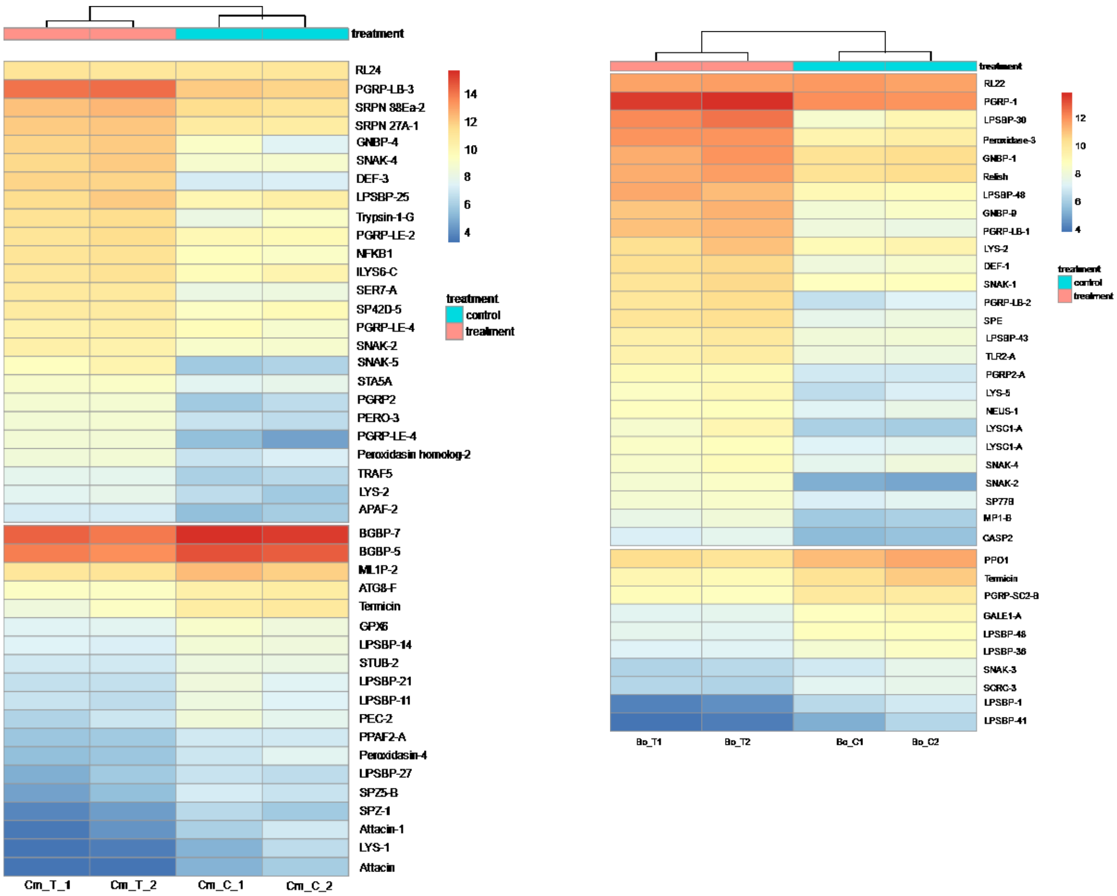


**Figure S4**. Differentially expressed immune genes after injection (left panel, subsocial cockroach, *C. meridianus*; right panel, solitary cockroach, *B. orientalis*).

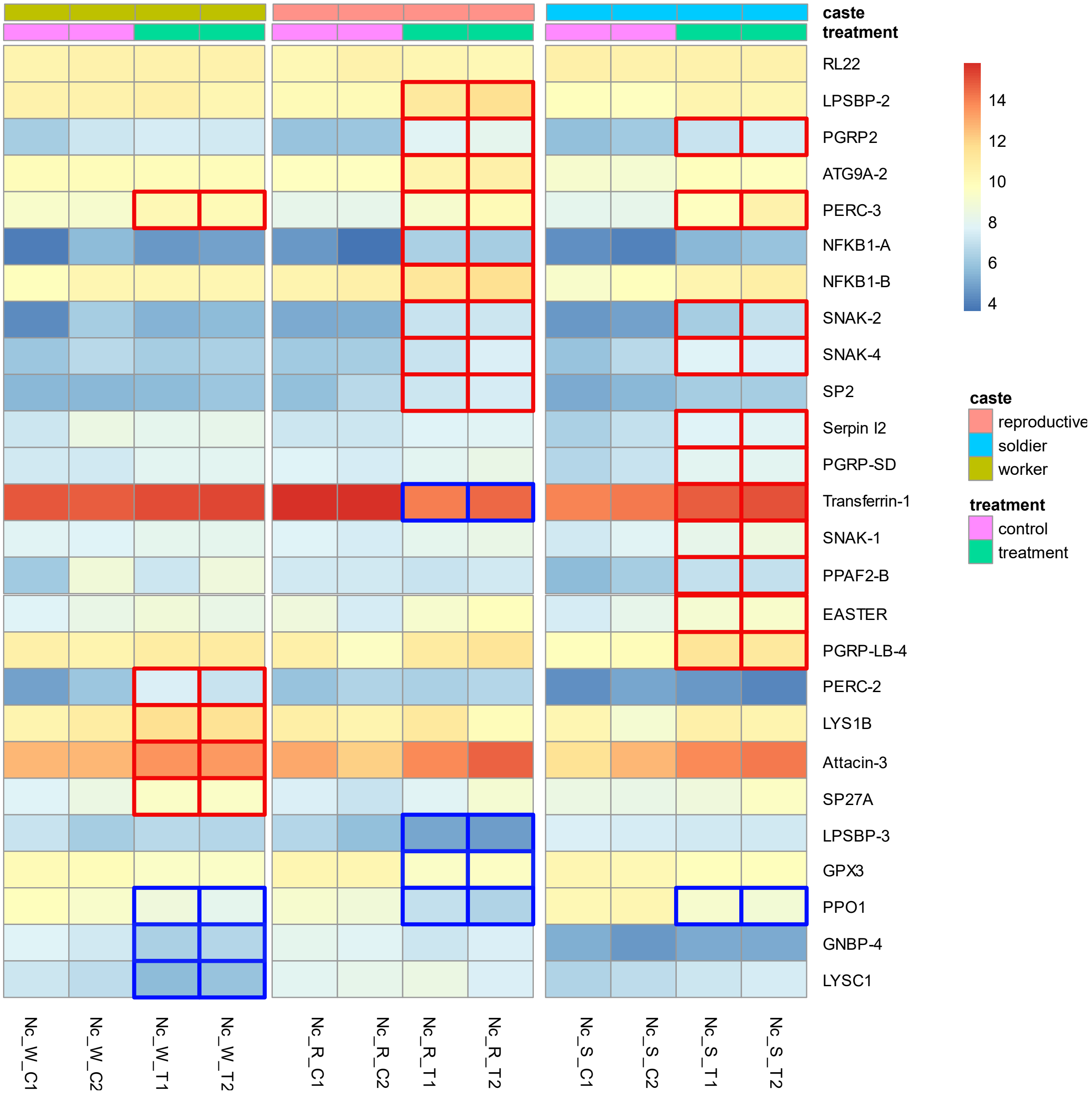


**Figure S5**. Differentially expressed immune genes in different castes after injection in individual experiment. Red squares indicate upregulated immune gene expression in caste after injected with heat killed microorganisms, and blue squares indicate downregulated immune gene expression in caste after injected with heat killed microorganisms.

| Species/caste | | Upregulated | | Downregulated | |
|---|---|---|---|---|---|
| | | non-immune | immune | non-immune | immune |
| *N. castaneus* | Reproductives | **211** | **9** | **384** | **4** |
| | Soldiers | **150** | **11** | **130** | **1** |
| | False Workers | **25** | **5** | **68** | **3** |
| | Total | **313** | **20** | **485** | **6** |
| *C. meridianus* | | **358** | **24** | **229** | **19** |
| *B. orientalis* | | **238** | **25** | **155** | **10** |

**Figure S6**. Summary numbers of differentially expressed genes (immune genes and non-immune genes) in 3 castes of the termite *N. castaneus*, subsocial cockroach *C. meridianus*, and solitary cockroach *B. orientalis* injected with heat-killed microbes versus Ringer's solution (**individual experiment**). The numbers of total differentially expressed genes are different from sum up of individual castes because of the shared genes among different castes.

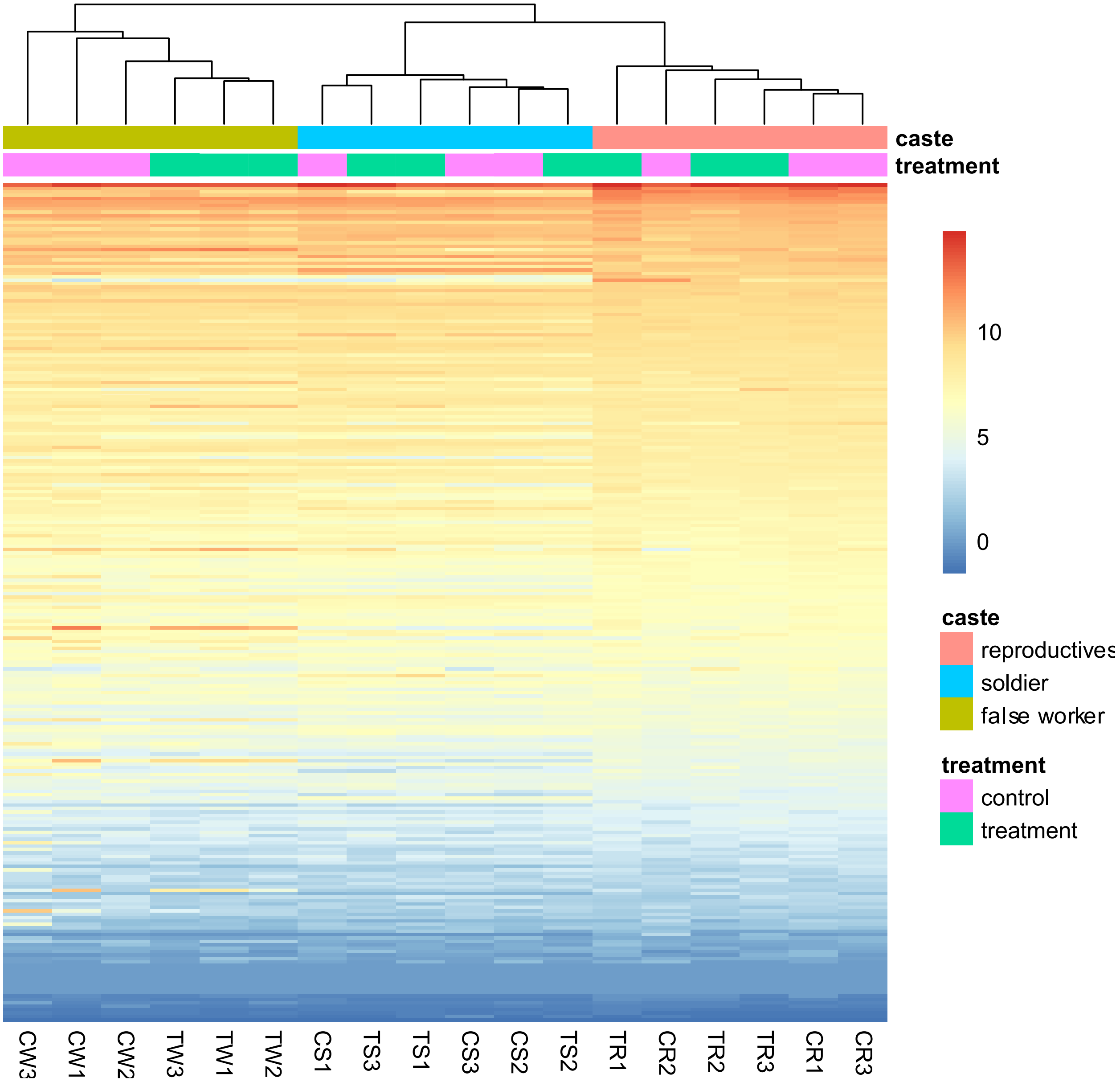


**Figure S7**. Heatmap of total identified immune gene expression in 3 *N. castaneus* termite castes following exposure to nestmate focal individuals injected with heat-killed microbes (treatment) or Ringer's solution (control) (**social experiment**). Library legend: CW1-3: false workers in control-exposed replicates, TW1-3: false workers in treatment-exposed replicates, CS1-3: soldiers in control-exposed replicates, TS1-3: soldiers in treatment-exposed replicates, CR1-3: reproductives in control-exposed replicates, TR1-3: reproductives in treatment-exposed replicates.

| Species/caste | | Upregulated | Downregulated |
| --- | --- | --- | --- |
| *Neotermes castaneus* | Reproductives | **1** | **1** |
| | Soldiers | **1** | **0** |
| | False Workers | **12** | **96** |
| *Blatta orientalis* | | **9** | **7** |

**Figure S8**. Number of differentially regulated genes in 3 castes of the termite *N. castaneus* or conspecifics of the cockroach *B. orientalis* following exposure to nestmate or conspecific focal individuals that had been injected with heat-killed microbes versus Ringer's solution (**social experiment**).

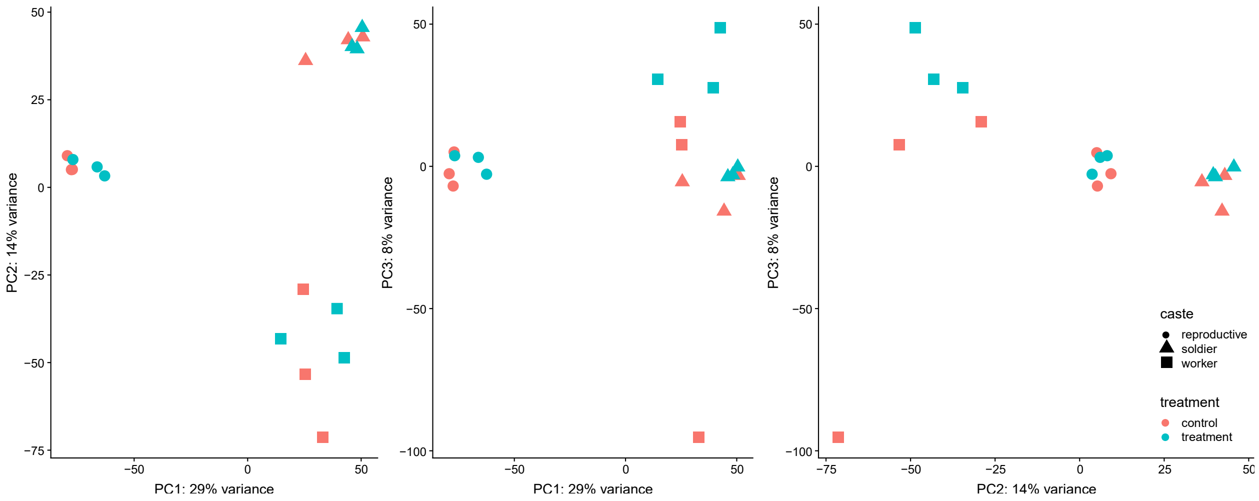


**Figure S9**. PCA analysis of immune gene expression in 3 different castes of the termite *N. castaneus* following exposure to nestmate focal individuals injected with heat-killed microbes (treatment, blue) versus Ringer's solution (control, red) (**social experiment**).

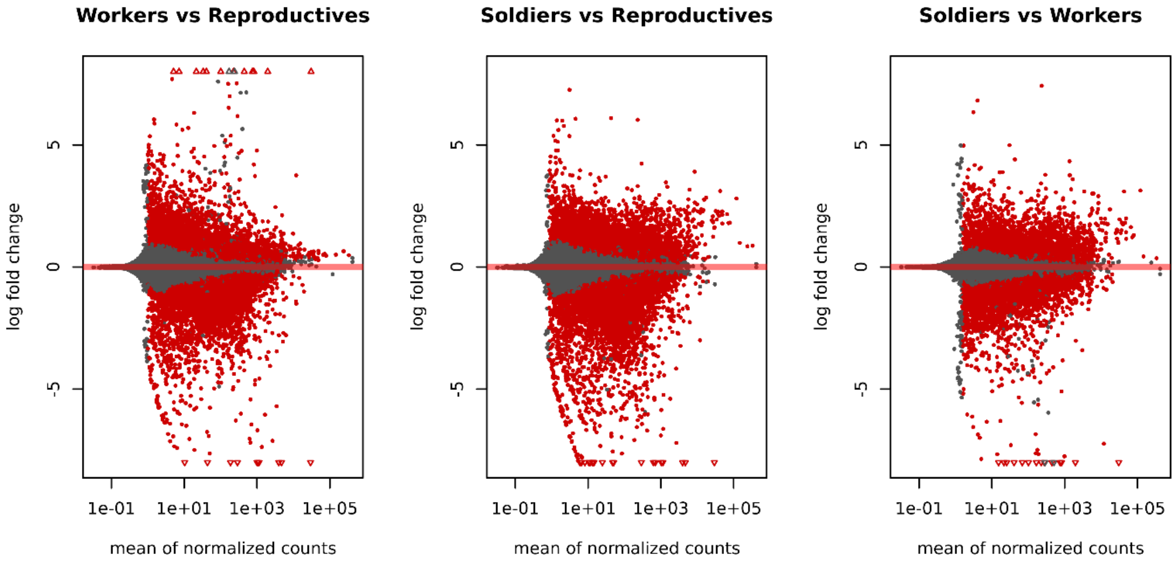


**Figure S10**. MA plots of total differential gene expression comparisons between the 3 different castes of the termite *N. castaneus*, regardless of social treatment exposure (combined treatment and control-exposure replicates from the **social experiment**). From left to right: false workers vs reproductives; soldiers vs reproductives; soldiers vs false workers.

| | Only soldiers | Soldier & False Worker | Only false workers |
|---|---|---|---|
| Reproductives-up | **601** | **770** | **165** |
| Reproductives-down | **421** | **28** | **436** |

| | Only reproductives | Reproductives & False Worker | Only false workers |
|---|---|---|---|
| Soldiers-up | **327** | **122** | **90** |
| Soldiers-down | **1275** | **96** | **321** |

| | Only reproductives | Reproductives & False Workers | Only soldiers |
|---|---|---|---|
| Workers-up | **275** | **189** | **238** |
| Workers-down | **912** | **23** | **189** |

**Figure S11**. Number of differentially expressed genes between the 3 different castes of the termite *N. castaneus*, regardless of social exposure (combined treatment- and control-exposure replicates from the **social experiment**) (|log2FoldChange| > 2 and p-value<0.01).
